# TRAF6 integrates innate immune signals to regulate glucose homeostasis via Parkin-dependent and-independent mitophagy

**DOI:** 10.1101/2025.01.31.635900

**Authors:** Elena Levi-D’Ancona, Emily M. Walker, Jie Zhu, Yamei Deng, Vaibhav Sidarala, Ava M. Stendahl, Emma C. Reck, Belle A. Henry-Kanarek, Anne C. Lietzke, Mabelle B. Pasmooij, Dre L. Hubers, Venkatesha Basrur, Sankar Ghosh, Linsey Stiles, Alexey I. Nesvizhskii, Orian S. Shirihai, Scott A. Soleimanpour

**Author notes:** Corresponding Author. Scott A. Soleimanpour, MD, 1000 Wall Street, Brehm Tower Room 6329, Ann Arbor, MI 48105, USA.

## Abstract

Activation of innate immune signaling occurs during the progression of immunometabolic diseases, including type 2 diabetes (T2D), yet the impact of innate immune signaling on glucose homeostasis is controversial. Here, we report that the E3 ubiquitin ligase TRAF6 integrates innate immune signals following diet-induced obesity to promote glucose homeostasis through the induction of mitophagy. Whereas TRAF6 was dispensable for glucose homeostasis and pancreatic β-cell function under basal conditions, TRAF6 was pivotal for insulin secretion, mitochondrial respiration, and increases in mitophagy following metabolic stress in both mouse and human islets. Indeed, TRAF6 was critical for the recruitment and function of machinery within both the ubiquitin-mediated (Parkin-dependent) and receptor-mediated (Parkin-independent) mitophagy pathways upon metabolic stress. Intriguingly, the effect of TRAF6 deficiency on glucose homeostasis and mitophagy was fully reversed by concomitant Parkin deficiency. Thus, our results implicate a role for TRAF6 in the cross-regulation of both ubiquitin-and receptor-mediated mitophagy through the restriction of Parkin. Together, we illustrate that β-cells engage innate immune signaling to adaptively respond to a diabetogenic environment.

## INTRODUCTION

Innate immune responses serve as the first line of defense against pathogens^1^. In immune cells, innate immune signaling pathways are regulated by toll-like receptors (TLRs), transmembrane receptors that recognize and respond to foreign ligands. Among the most studied innate immune signaling receptors is TLR4, which primarily recognizes components of gram-negative bacteria, but also mediates signaling as a sensor for free fatty acids^2,3^. TLR4 activation leads to a complex signal transduction network via the signaling adaptors MYD88 and TRIF, which interact with the E3 ubiquitin ligase TNF receptor associated factor 6 (TRAF6), a nexus for host defense against microbial pathogens^2^. While the importance of TLRs is well-known in the immune system, the importance of innate immune signaling in non-immune cells of metabolic tissues, where engagement with microbial pathogens is not as common, is less clear.

TLRs are expressed in numerous cell types associated with systemic glucose homeostasis, including adipocytes, hepatocytes and insulin-producing pancreatic β-cells^6–8^. TLR expression and signaling increases during obesity and the subsequent development of metabolic syndrome to drive systemic inflammation^3,7,9^. Furthermore, *TRAF6* expression is elevated in peripheral blood mononuclear cells of individuals with type 2 diabetes (T2D)^10^, and *TRAF6* polymorphisms are associated with diabetic complications^11,12^. Exposure of mouse or human pancreatic islets to TLR4 ligands, such as endotoxin, impairs β-cell function and viability through intra-islet cytokine production and inhibition of insulin gene expression^13–17^. While classical TLR4 ligands can be detrimental to β-cells, the absence or inhibition of TLR4 signaling components have had contradictory effects. Whole-body deletion of both *Tlr2* and *Tlr4* was previously reported to improve β-cell proliferation following diet-induced obesity^8^, and inhibition of TLR4 prevented diabetes onset in autoimmune NOD mice^18^, demonstrating that loss of TLR4 signaling is favorable for β-cell health and function during diabetes. In contrast, mice null for either *Myd88* or *Trif* develop impaired β-cell function^19,20^. The paradoxical findings of these previous studies may relate to their reliance on whole-body knockout models, suggesting the role of TLR4-mediated innate immune signaling in β-cells remains to be clarified.

In immune cells, mitochondria are an integral part of innate immune signaling. Immune TLR signaling regulates oxidative metabolism, rewires metabolic pathways to generate ROS for microbial killing, releases mitochondrial nucleic acids to amplify local and systemic innate immune signals, and engages mitochondrial docking sites for signaling cascades by the viral sensing machinery^21^. In β-cells, mitochondria provide the energy necessary to support insulin biosynthesis and secretion following glucose stimulation^22–25^. β-cell mitochondria are highly susceptible to inflammatory or metabolic stress that occurs during the development of diabetes^26,27^, and mitochondrial defects have been observed to precede the development of T2D^28–31^. To preserve mitochondrial health, insulin release, and glycemic control following metabolic stress, β-cells engage mitophagy (*i*.*e*., the selective clearance of mitochondria by autophagy) through ubiquitin-or receptor-mediated quality control pathways that act dependently or independently of the E3 ligase Parkin, respectively^32,33^. Moreover, we recently observed that β-cells of human islet donors with T2D have defects in mitophagy^34^, and loss of function variants of *CALCOCO2*, which encodes a ubiquitin receptor necessary for mitophagy, have been linked to T2D^35^. However, the effect of innate immune signaling on β-cell mitochondrial respiration or quality control is unknown.

Here, we show that β-cell innate immune signaling via TRAF6 is important for glucose homeostasis during metabolic stress via the regulation of mitochondrial quality control. We found that β-cells engage innate immune pathways following fatty acid exposure that models the lipotoxicity observed in T2D. Further, we found that impairment of TRAF6 function *in vivo* elicits glucose intolerance and reduced insulin secretion during high fat diet-induced metabolic stress. Using quantitative proteomics, high-resolution imaging, biochemical assays, and transcriptomic profiling, we demonstrate that TRAF6-deficiency leads to an accumulation of dysfunctional mitochondria due to impaired recruitment of components of the ubiquitin-and receptor-mediated mitophagy machinery. We also found that TRAF6 is essential for mitophagy and insulin secretion following metabolic stress in human islets. Notably, impaired glucose homeostasis and defective mitophagy in TRAF6-deficient β-cells following metabolic stress is fully reversed by Parkin-deficiency, implicating a novel role for TRAF6 in the cross-regulation of ubiquitin-and receptor-mediated mitophagy through restriction of Parkin. Together, our results illustrate that β-cells engage innate immune signaling to promote mitophagy as a vital adaptive response to metabolic stress.

## RESULTS

### Palmitate activates innate immune signaling responses in pancreatic β-cells

We hypothesized that β-cells employ innate immune signaling in response to diabetogenic stimuli based on studies exploring TLR signaling components in β-cells^8,19,20^. To test this notion, we examined a previously generated RNA-seq dataset from islets isolated from mice fed high fat diet (HFD) or regular fat diet (RFD) for 12 weeks^36^. We observed over 770 differentially expressed genes (DEGs) following diet-induced obesity (DIO), including expected changes in lipid and fatty acid metabolism follow HFD-feeding (**Figure S1A-D**). We also observed increases in type I interferon-responsive pathways (**Figure S1A-D**), as well as expression of chemokines (*i*.*e*., *Ccl5* and *Cxcl10*) and other key response genes activated by innate immune signaling (*i*.*e*., *Oas2*, *Irf7*, and *Isg15*) (**Figure S1A**). These increases in innate immune signaling targets and lipid metabolic pathways are consistent with fatty acid-triggered TLR signaling, and they supported the concept that DIO could induce innate immune signaling in β-cells.

To isolate the effects of metabolic stress in β-cells without interference from local immune cells, we next performed a screen in Min6 β-cells exposed to metabolic stress conditions in culture to identify which members of the innate immune signaling pathways downstream of TLR4 were activated. We transfected Min6 cells with transcriptional reporter plasmids for pathways responsive to innate immune signaling^37,38^ (**Table S1**), including NF-κB (NF-κB and RELB), MAPK, JAK/STAT (STAT1 and STAT3), interferon signaling (IRF3, IFNB1, Type I Interferon and Type II Interferon), or stress response pathways (p53, ATF4, C/EBPβ and TFEB). These reporters were then examined in β-cells in the context of two exposures of metabolic stress; namely, lipopolysaccharide (LPS), a classical TLR4 ligand that is known to be elevated during obesity due to gut dysbiosis^4,39^, or the saturated fatty acid palmitate. Palmitate both emulates the lipotoxicity observed in T2D and induces innate immune/TLR4 signaling in immune cells^38,40^, but its impact on β-cell innate immune signaling was previously unknown. We found LPS and palmitate elicited similar patterns of reporter activity in β-cells, albeit to differing magnitudes (**Figure 1A**).

**Figure 1.**
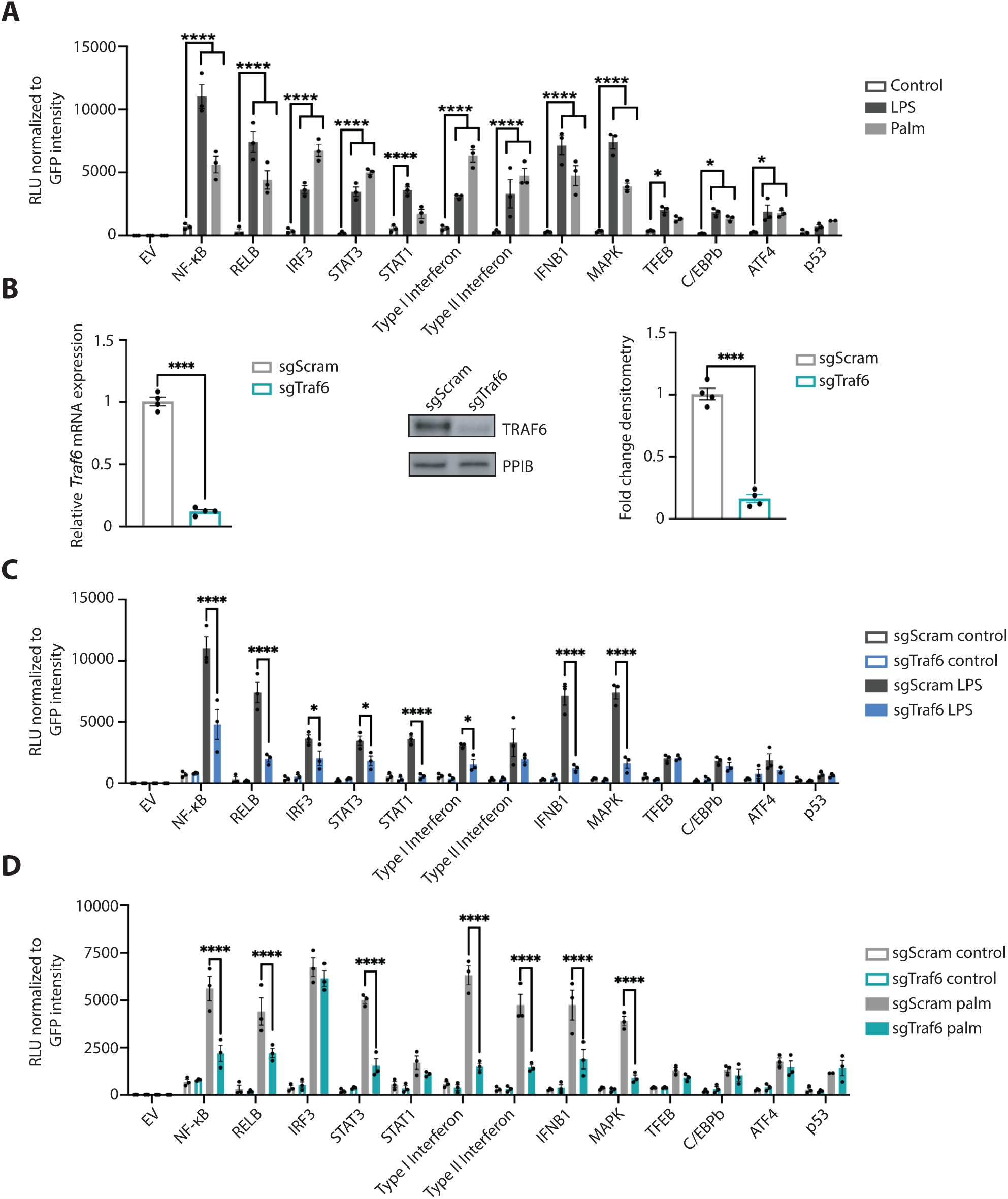
Palmitate activates innate immune pathways in β-cells. **(A)** Luciferase activity, normalized to GFP intensity, of Min6 β-cells transfected with empty vector (EV), NFκB, RELB, IRF3, STAT3, STAT1, Type I Interferon, Type II Interferon, IFNB1, MAPK, TFEB, C/EBPβ, ATF4, and p53 reporter plasmids following exposure to vehicle (DMSO + BSA), LPS (5 nM for 2 h), or palmitate (0.5 mM for 48 h). *n* = 3/group. \*\**P* < 0.01, \*\*\**P* < 0.001, \*\*\*\**P* < 0.0001 by 2-way ANOVA. **(B)** qPCR analysis of *Traf6*, normalized to *Hprt* expression, (left) and TRAF6 protein expression by western blot (WB), quantified by densitometry (right), in CRISPR-generated sgScram or sgTraf6 Min6 β-cells. Cyclophilin B (PPIB) served as a loading control. *n* = 4/group. \*\*\*\**P* < 0.0001 by Student’s unpaired t-test. (**C)** Luciferase activity, normalized to GFP intensity, of sgScram and sgTraf6 Min6 cells transfected with reporter plasmids following exposure to vehicle or 5 nM LPS for 2 h. *n* = 3/group. \*\*\**P* < 0.001, \*\*\*\**P* < 0.0001 by 2-way ANOVA. **(D)** Luciferase activity, normalized to GFP intensity, of sgScram and sgTraf6 Min6 cells transfected with reporter plasmids following exposure to vehicle or 0.5 mM palmitate for 48 h. *n* = 3/group. \*\*\*\**P* < 0.0001 by 2-way ANOVA.

To determine if the effects of palmitate in β-cells were mediated by innate immune signaling, we generated Min6 β-cells deficient in TRAF6, an E3 ubiquitin ligase that acts as an innate immune signaling hub^41^, by CRISPR-Cas9 gene editing. We also generated Min6 β-cells transduced with a scrambled guide RNA control (**Figures 1B-C**, hereafter known as sgTraf6 and sgScram, respectively). Immune-responsive reporter activity was significantly lower in the TRAF6-deficient cells exposed to LPS compared to the LPS-exposed sgScram cells, confirming TRAF6 function within the innate immune signaling pathway in β-cells (**Figure 1C**). The effects of palmitate were similarly blunted in TRAF6-deficient cells (**Figure 1D**). We similarly found that LPS and palmitate exposure each induced expression of the NF-κB target genes *inducible nitric oxide synthase* (*iNOS)* and *super oxide dismutase 2* (*Sod2*)^42^, which were also TRAF6-dependent (**Figure S1E-F**). Taken together, these data suggest that innate immune signaling is induced in β-cells in the setting of metabolic stress.

### TRAF6 promotes β-cell functional compensation in response to diet-induced obesity

To clarify the importance of innate immune signaling in β-cells *in vivo*, we generated mice bearing β-cell-specific deletion of *Traf6* (*Traf6^loxP/loxP^;Ins1^Cre^,* hereafter known as Traf6^Δβ^; **Figure 2A**).

**Figure 2.**
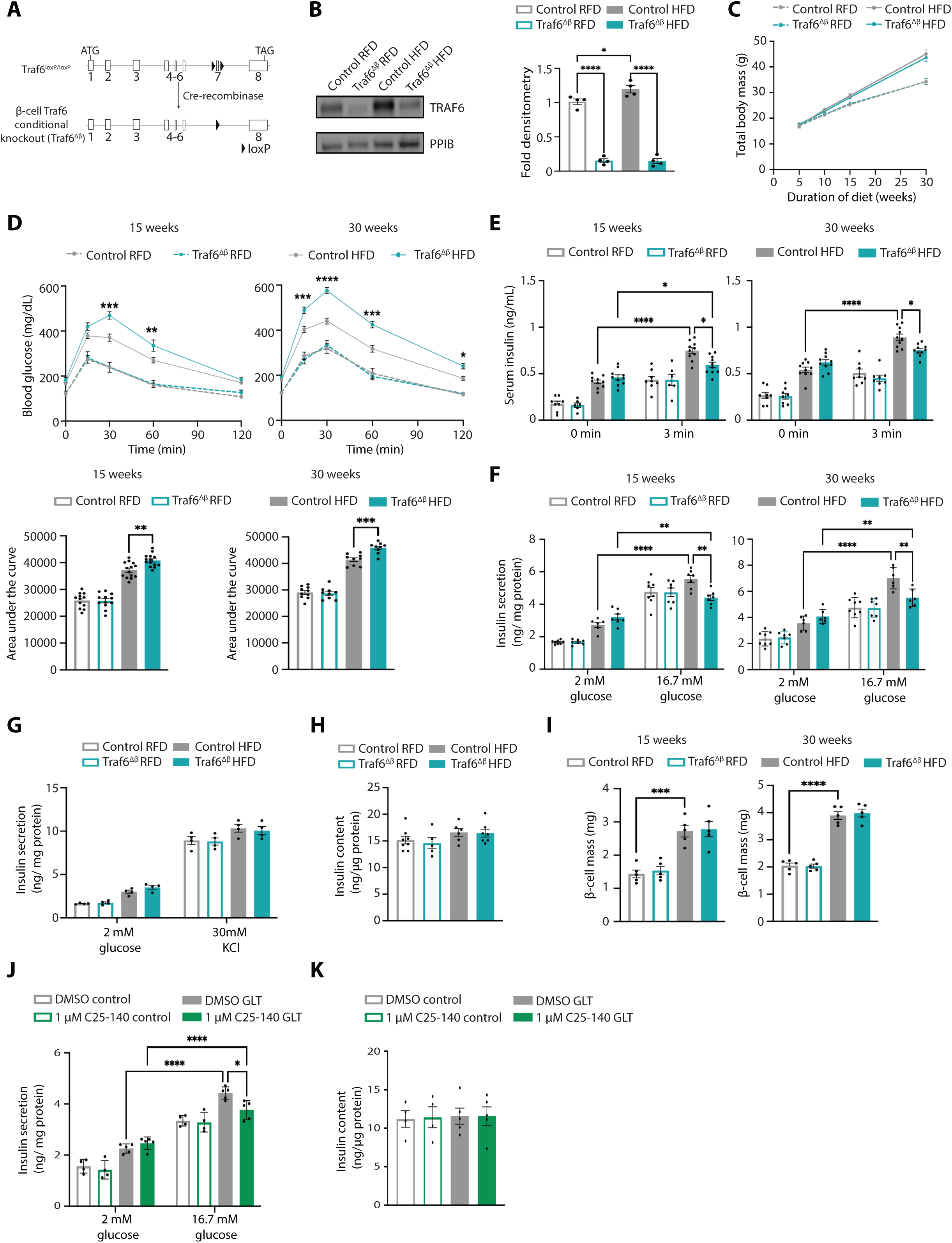
TRAF6 preserves insulin secretion following metabolic stress in mouse and human β-cells. **(A)** A schematic diagram illustrating Cre-mediated recombination at the *Traf6* locus. **(B)** Protein expression and densitometry quantification (relative to Control RFD) of TRAF6 by WB in islets isolated from control or Traf6^Δβ^ mice fed regular fat diet (RFD) or high fat diet (HFD) for 15 weeks. PPIB served as a loading control. *n* = 4/group. \**P* < 0.05, \*\*\*\**P* < 0.0001 by 1-way ANOVA. **(C)** Total body mass of male control or Traf6^Δβ^ mice following 30 weeks of RFD or HFD. *n* = 7-10/group. **(D)** Blood glucose concentrations measured during intraperitoneal glucose tolerance test (IPGTT; top) and area under the curve (AUC; bottom) of male control and Traf6^Δβ^ mice fed either RFD (dashed lines) or HFD (solid lines) for 15 weeks (left) or 30 weeks (right). *n* = 8-12/group. \*\**P* < 0.01, \*\*\**P* < 0.001, \*\*\*\**P* < 0.0001 by 2-way ANOVA (IPGTT) or 1-way ANOVA (AUC). **(E)** Serum insulin levels measured during *in vivo* glucose-stimulated insulin release assays in male control and Traf6^Δβ^ mice fed RFD or HFD for 15 weeks (left) or 30 weeks (right). *n* = 8-10/group. \**P* < 0.05, \*\*\*\**P* < 0.0001 by 1-way ANOVA. **(F)** Glucose-stimulated insulin secretion from islets isolated from control or Traf6^Δβ^ mice fed RFD or HFD for 15 weeks (left) or 30 weeks (right). *n* = 6-8/group. \*\**P* < 0.01, \*\*\*\**P* < 0.0001 by 2-way ANOVA. **(G)** KCl-stimulated insulin secretion from islets isolated from control or Traf6^Δβ^ mice fed RFD or HFD for 15 weeks. *n* = 4/group. **(H)** Insulin content in islets from control or Traf6^Δβ^ mice fed 15-week RFD or HFD. *n* = 5-8/group. **(I)** Pancreatic β-cell mass from control or Traf6^Δβ^ mice fed RFD or HFD for 15 weeks (left) or 30 weeks (right). *n* = 5/group. **(J)** Glucose-stimulated insulin secretion following static incubations in 2 mM and 16.7 mM glucose, performed in human islets treated with DMSO or 1 μM C25-140 (TRAF6 inhibitor) for 24 h and exposed to glucolipotoxicity (GLT; 25 mM glucose + 0.5 mM palmitate) or control (5 mM glucose + BSA) for 48 h. *n* = 4-5/group. \**P* < 0.05, \*\*\*\**P* < 0.0001 by 2-way ANOVA. **(K)** Insulin content in human islets treated with DMSO or 1 μM C25-140 for 24 h and exposed to GLT or control for 48 h. *n* = 4-5/group.

Traf6^Δβ^ mice exhibited an ∼80% reduction in TRAF6 protein levels in islets compared to littermate controls with no changes in body weight (**Figure 2B-C**). We observed that TRAF6 was dispensable for glucose tolerance following both 15-week and 30-week RFD (**Figure 2D**).

To determine the importance of innate immune signaling in β-cell responses to metabolic stress *in vivo*, we fed mice a HFD for up to 30 weeks beginning at weaning. Notably, Traf6^Δβ^ mice and littermate controls exhibited a nearly identical trajectory of weight gain following HFD feeding (a lack of difference in weight gain between the two groups was also seen when they were both fed a RFD) (**Figure 2C**). Interestingly, expression of TRAF6 protein (**Figure 2B**), but not mRNA (**Figure S1A**), was elevated in islets of control mice following DIO.

Given our observations of innate immune signaling engagement and stress responses by palmitate exposure that were dependent on TRAF6 (**Figure 1**), we speculated that TRAF6-deficient β-cells would exhibit improved metabolic function *in vivo* following HFD. However, we found that male Traf6^Δβ^ mice were not protected from DIO-related metabolic dysfunction as they exhibited progressive glucose intolerance following HFD feeding (**Figure 2D, S2A**). We also observed no differences in insulin tolerance between genotypes following either RFD or HFD feeding, suggesting that alterations in insulin sensitivity were not responsible for the glucose intolerance observed following TRAF6-deficiency (**Figure S2B**). Rather, HFD-fed male Traf6^Δβ^ mice exhibited less insulin release *in vivo* after a glucose challenge compared to littermate controls (**Figure 2E**). We also observed reduced glucose-stimulated insulin secretion (GSIS) in isolated islets of HFD-fed TRAF6-deficient mice compared to islets from HFD-fed control mice (**Figure 2F**), yet both *in vivo* and *ex vivo* insulin release were unchanged in RFD-fed TRAF6-deficient mice (**Figure 2E-F**). There were no differences in insulin secretion following KCl stimulation of isolated Traf6^Δβ^ islets (**Figure 2G**). TRAF6-deficient mice also displayed no changes in insulin content or β-cell mass compared to littermate controls and appropriate expansion of β-cell mass in response to HFD (**Figure 2H-I**). We also observed similar impairments in glucose homeostasis and insulin release without differences in body weight or insulin sensitivity following DIO in Traf6^Δβ^ females (**Figure S3A-D**). Taken together, these data suggest that reductions in glucose tolerance and GSIS in HFD-fed Traf6^Δβ^ mice were not due to defects in insulin content, β-cell number, or insulin granule exocytosis.

To next determine if TRAF6 was also necessary for β-cell functional compensation during metabolic stress in human islets, we treated primary human islets with the TRAF6 inhibitor C25-140, which disrupts the interaction between TRAF6 and the E2 conjugating enzyme UBC13^43^. We also exposed human islets to glucolipotoxic conditions (GLT; 25 mM glucose and 0.5 mM palmitate) to model the diabetogenic metabolic stress of hyperglycemia and hyperlipidemia observed during T2D^44^. We first confirmed that C25-140 inhibits TRAF6 activity by observing a dose-dependent decrease in NF-κB target gene activation following exposure to a pro-inflammatory cytokine cocktail (**Figure S4**). Similar to our observations in HFD-fed Traf6^Δβ^ islets, we found that inhibiting TRAF6 function reduced GSIS in human islets following GLT (**Figure 2J**), which was not due to a reduction in insulin content (**Figure 2K**). Moreover, TRAF6 was dispensable for GSIS in human islets in the absence of metabolic stress (**Figure 2J**), again replicating our observations in RFD-fed Traf6^Δβ^ mice. Taken together, these results demonstrate that TRAF6 is vital for adaptation to metabolic stress by preserving GSIS in both mouse and human β-cells.

### TRAF6-deficiency results in an accumulation of dysfunctional mitochondria following HFD

To understand how TRAF6 maintains β-cell insulin secretion during metabolic stress, we evaluated whether TRAF6 regulates mitochondrial function in β-cells given the importance of mitochondria for insulin release and TLR signaling^21,24^. We measured mitochondrial respiration in isolated islets from Traf6^Δβ^ mice and littermate controls following 15-weeks of RFD or HFD feeding. We observed significantly lower glucose-stimulated oxygen consumption in islets isolated from HFD-fed Traf6^Δβ^ mice compared to HFD-fed control mice (**Figure 3A**). Additionally, we saw significant reductions in glucose-stimulated ATP/ADP ratio in islets from HFD-fed Traf6^Δβ^ mice (**Figure 3B)**. However, there were no differences in mitochondrial function between the genotypes following RFD feeding (**Figure 3A-B**).

**Figure 3.**
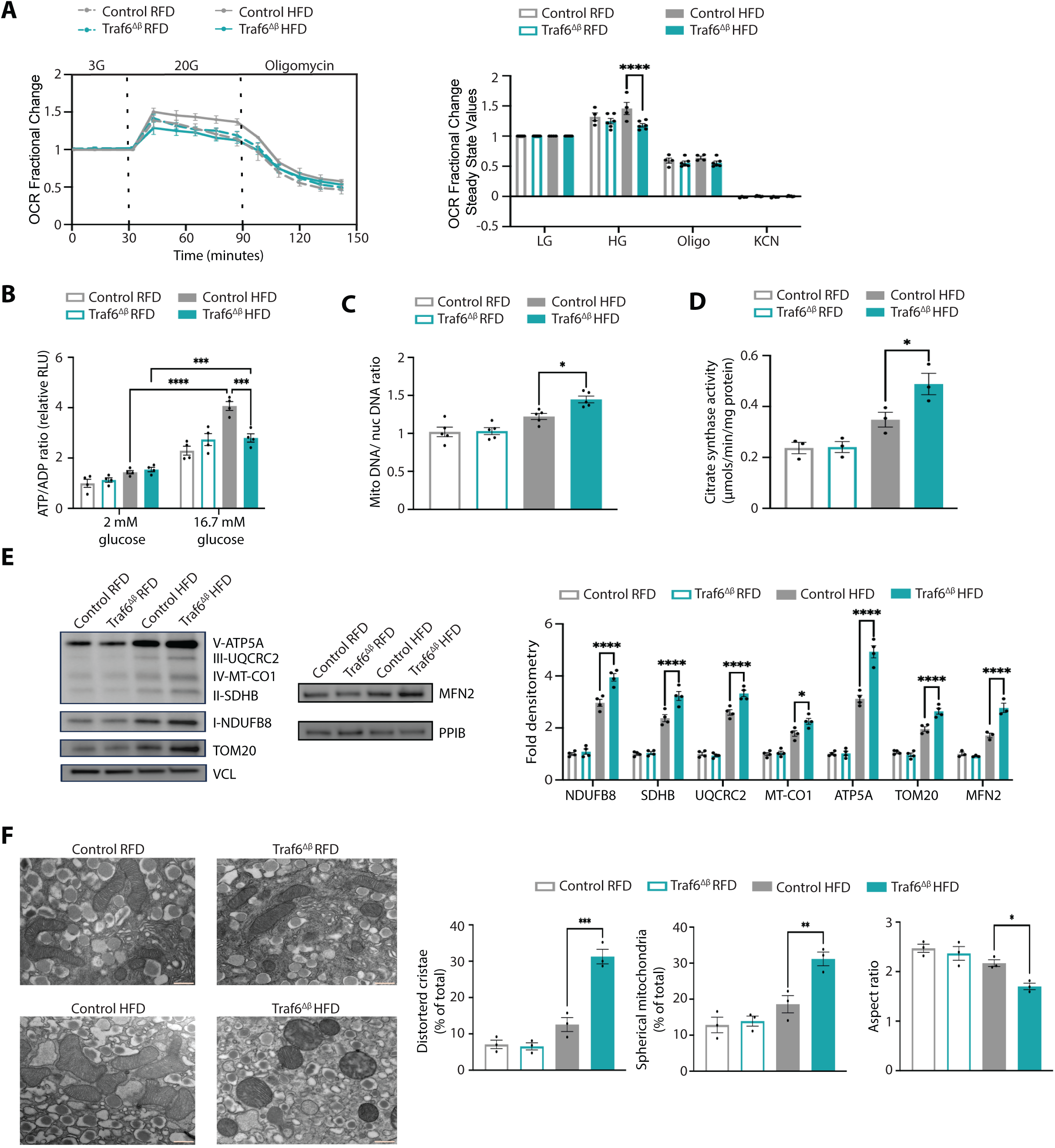
TRAF6-deficiency results in accumulation of dysfunctional mitochondria following DIO. **(A)** Oxygen consumption rate (OCR) fractional change (left) and steady state quantification (right) following exposure to 3 mM glucose, 20 mM glucose, 10 μM oligomycin, and 3 mM KCN in isolated islets from control or Traf6^Δβ^ mice fed 15-week RFD or HFD via Barofuse respirometry. *n* = 4-6/group. \*\*\*\**P* < 0.0001 by 2-way ANOVA. **(B)** ATP/ADP ratio measured in isolated islets from control or Traf6^Δβ^ mice fed RFD or HFD for 15 weeks. *n* = 4/group. \*\*\**P* < 0.001, \*\*\*\**P* < 0.0001 by 2-way ANOVA. **(C)** mtDNA to nuclear DNA ratio in isolated islets from control or Traf6^Δβ^ mice fed RFD or HFD for 15 weeks by qPCR. *n* = 5/group. \**P* < 0.05 by 1-way ANOVA. **(D)** Citrate synthase activity in isolated islets from control or Traf6^Δβ^ mice fed RFD or HFD for 15 weeks. *n* = 3/group. * *P* < 0.05 by 1-way ANOVA. **(E)** WB for OXPHOS subunits, TOM20, and MFN2 with densitometry (relative to Control RFD) in islets from control or Traf6^Δβ^ mice fed RFD or HFD for 15 weeks. Vinculin (VCL) or PPIB served as loading controls. *n* = 4/group. \**P* < 0.05, \*\*\*\**P* < 0.0001 by 2-way ANOVA. **(F)** Representative transmission electron microscopy (TEM) image of β-cells from control or Traf6^Δβ^ mice fed RFD or HFD for 15 weeks with quantification of mitochondrial morphology (∼100 independent mitochondria scored/animal). *n* = 3/group. \**P* < 0.05, \*\**P* < 0.01, \*\*\**P* < 0.001 by 1-way ANOVA. Scale bars: 500 nM.

To clarify the etiology of mitochondrial respiratory dysfunction following TRAF6-deficiency, we next examined mitochondrial mass and ultrastructure. Several measures indicated increased mitochondrial mass in islets from HFD-fed Traf6^Δβ^ mice, including increased mtDNA content, citrate synthase activity, and expression of OXPHOS subunits, TOM20, and MFN2 (**Figure 3C-E**), indicating reduced mitochondrial respiration in TRAF6-deficient β-cells was not due to fewer mitochondria. Transmission electron microscopy (TEM) demonstrated that TRAF6-deficient β-cells from HFD-fed mice exhibited abnormal mitochondrial morphology compared to littermate controls as determined by increases in mitochondria with distorted cristae and spherical appearance. In addition, TRAF6-deficient β-cells from HFD-fed mice showed reductions in mitochondrial aspect ratio, suggestive of mitochondrial fragmentation (**Figure 3F**)^45^. Together, these results indicated that TRAF6-deficient β-cells accumulate dysfunctional mitochondria following DIO.

### TRAF6 promotes mitophagy in β-cells following diet-induced obesity

Mitochondrial mass is regulated by a balance of mitochondrial biogenesis and clearance^46^. Thus, we next assessed whether the accumulation of mitochondria in islets from HFD-fed Traf6^Δβ^ mice were driven by changes in biogenesis and/or mitophagy. We observed no differences in the expression of key regulators of mitochondrial biogenesis following TRAF6-deficiency^46–48^ (**Figure 4A)**. Next, we quantified β-cell mitophagy via flow cytometry utilizing the pH sensitive MtPhagy dye to identify mitochondria within the acidic lysosome compartment, with and without exposure to the potassium ionophore valinomycin to elicit mitophagic flux, as previously described^49^. In control mice, exposure to HFD led to greater β-cell mitophagic flux, but this effect was blunted in β-cells from Traf6^Δβ^ mice **(Figure 4B**). We observed no differences in β-cell mitophagic flux following TRAF6-deficiency in RFD-fed mice, nor did we observe differences in mitophagic flux following TRAF6-deficiency in non β-cells following either RFD or HFD (**Figure 4B**). As an orthogonal approach, we assessed mitophagic flux in isolated islets from RFD-or HFD-fed mt-Keima transgenic mice, which express a mitochondrial-targeted dual excitation reporter that shifts its excitation spectra in a pH-specific manner^50^. As expected, mitophagic flux was increased in β-cells from HFD-fed mt-Keima mice, but this effect was lost upon inhibition of TRAF6 with C25-140, phenocopying our results in β-cells from HFD-fed Traf6^Δβ^ mice (**Figure 4C)**. Pharmacologic inhibition of TRAF6 in human islets also impaired β-cell mitophagic flux following GLT, whereas TRAF6 was dispensable for human β-cell mitophagic flux in the absence of metabolic stress (**Figure 4D**), similar to our observations in RFD-fed mice.

**Figure 4.**
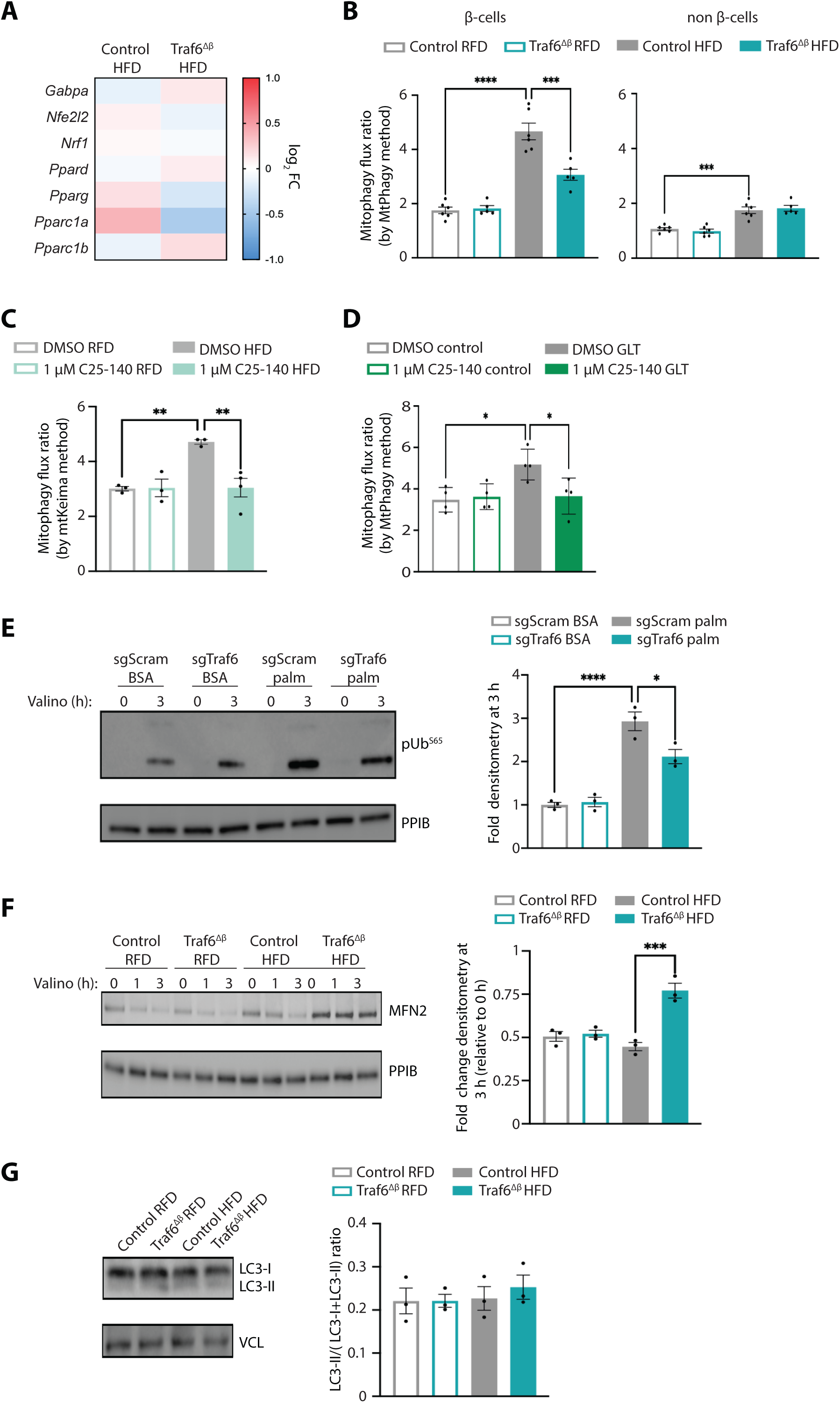
TRAF6 promotes mitophagy in mouse and human β-cells following metabolic stress. **(A)** Differential RNA expression heatmap of selected mitochondrial biogenesis genes in isolated islets from control or Traf6^Δβ^ mice fed HFD for 15 weeks. *n* = 3-4/group. **(B)** Flow cytometry quantification of mitophagy flux by MtPhagy dye approach in β-cells from control or Traf6^Δβ^ mice fed RFD or HFD for 15 weeks. *n* = 5-6/group. \*\*\**P* < 0.001 by 1-way ANOVA. **(C)** Flow cytometry quantification of mitophagy flux in β-cells from mt-Keima mice fed RFD or HFD for 15 weeks, followed by ex vivo exposure to DMSO or 1 μM C25-140 for 24 h. *n* = 3-4/group. \*\**P* < 0.01 by 1-way ANOVA. **(D)** Flow cytometry quantification of mitophagy flux by MtPhagy dye approach in human β-cells following exposure to DMSO or 1 μM C25-140 for 24 h and exposed to GLT or control for 48 h. *n* = 4/group. \**P* < 0.05 by 1-way ANOVA. **(E)** WB with densitometry of phosphorylation of ubiquitin at serine 65 (pUb^S65^) in sgScram and sgTraf6 Min6 cells following exposure to BSA or 0.5 mM palmitate for 48 h and 250 nM valinomycin for the final 3 h. PPIB served as a loading control. *n* = 3/group. \**P* < 0.05 by 1-way ANOVA. **(F)** WB with densitometry of MFN2 in isolated islets from control or Traf6^Δβ^ mice fed RFD or HFD for 15 weeks following 250 nM valinomycin exposure for up to 3 h. Densitometry represents relative change in MFN2 protein at 3 h following valinomycin exposure relative to 0 h baseline. PPIB served as a loading control. *n* = 3/group. \*\*\**P* <0.001 by 1-way ANOVA. **(G)** WB with densitometry of LC3-II expression (ratio of LC3-II/(LC3-I+LC3-II)) in isolated islets from control or Traf6^Δβ^ mice fed RFD or HFD for 15 weeks. VCL served as an additional independent loading control. *n* = 3/group.

To further clarify these defects in mitophagy, we assessed phosphorylation of ubiquitin at serine 65 (phospho-Ser65-Ub), a key early event in ubiquitin-mediated/Parkin-dependent mitophagy^51,52^. Following palmitate exposure in Min6 β-cells, valinomycin stimulated a robust increase in phospho-Ser65-Ub, which was markedly lower in TRAF6-deficient Min6 cells (**Figure 4E**). We next measured turnover of the outer mitochondrial membrane protein mitofusin 2 (MFN2) following mitochondrial damage, a key target for ubiquitin-mediated clearance in Parkin-dependent mitophagy^53^. As expected, MFN2 levels were reduced following valinomycin exposure in islets from both RFD-and HFD-fed control mice (**Figure 4F**). However, MFN2 turnover following valinomycin was lower in islets from HFD-fed Traf6^Δβ^ mice compared to HFD-fed control mice, again indicative of impaired Parkin-dependent mitophagy (**Figure 4F**). There were no differences in LC3-II expression in Traf6^Δβ^ islets compared to littermate controls, suggesting that these mitochondrial defects were specific to mitophagy rather than a general defect in macroautophagy (**Figure 4G**). Taken together, these results indicated that TRAF6 regulates β-cell mitochondrial function during metabolic stress in both human and mouse islets via control of mitophagy.

To gain insight into how TRAF6 maintains β-cell mitochondrial quality control following DIO, we performed RNA-seq on islets from HFD-fed TRAF6-deficient mice and littermate controls. We observed 445 DEGs between HFD-fed Traf6^Δβ^ and control mice (**Figure S5A**). Pathway analysis of DEGs following TRAF6-deficiency highlighted changes in pathways such as cell migration and cell-matrix adhesion, yet did not identify known regulators of mitochondrial quality control (**Figure S5B**).

### ECSIT does not contribute to HFD-induced β-cell dysfunction following TRAF6-deficiency

As our transcriptomic studies did not reveal a clear mechanism for impaired mitophagy following TRAF6-deficiency, we next examined potential post-transcriptional regulators of mitochondrial function. We first examined established interacting partners of TRAF6 important for mitochondrial function and innate immune signaling. Evolutionarily Conserved Signaling Intermediate in Toll pathways (ECSIT) is a signaling adaptor that interacts with TRAF6 downstream of TLR activation, preventing excess inflammation in macrophages by operating as a negative feedback regulator^54^. ECSIT also regulates mitochondrial complex I stability, mitochondrial ROS, and mitophagy in neuroglioma cells and macrophages^55–57^. We noted lower complex I function and higher ROS levels in islets from HFD-fed Traf6^Δβ^ mice compared to HFD-fed littermate controls (**Figure S6A-B**), supporting the hypothesis that TRAF6 could act in coordination with ECSIT to regulate β-cell mitochondrial function following metabolic stress.

To establish whether TRAF6 and ECSIT interact in β-cells following metabolic stress, we immunoprecipitated TRAF6 in Min6 β-cells. We found that TRAF6 interacted with ECSIT at baseline and that this interaction increased following palmitate exposure (**Figure S6C**). ECSIT expression was elevated in islets following HFD and was further increased in the setting of TRAF6-deficiency (**Figure S6D**). Given that ECSIT resides in both the cytoplasm and mitochondria^55^, we assessed whether the increases in ECSIT following metabolic stress and TRAF6-deficiency were localized to the mitochondria. Indeed, biochemical subcellular fractionation of Min6 cells showed that both TRAF6 and ECSIT exhibited greater mitochondrial localization following palmitate exposure (**Figure S6E**). ECSIT localization to mitochondria was further enhanced in TRAF6-deficient cells following palmitate exposure (**Figure S6E**).

To explore if elevations in ECSIT levels might contribute to β-cell dysfunction in Traf6^Δβ^ mice, we crossed Traf6^Δβ^ mice (or *Ins1*-Cre controls) with mice bearing an *Ecsit* conditional allele (*Ecsit*^loxP^). In order to reduce ECSIT levels but not elicit metabolic dysfunction, we generated mice haploinsufficient for *Ecsit* in β-cells (*Ecsit*^loxP/+^; *Ins1*^Cre^, hereafter known as ECSIT^β-het^). ECSIT^β-het^ mice exhibited the expected reductions in ECSIT expression and did not develop defects in glucose homeostasis, GSIS, or insulin content during either RFD or HFD feeding (**Figures S7A-J)**. Islets from Traf6^Δβ^/ECSIT^β-het^ mice exposed to HFD displayed lower levels of both TRAF6 and ECSIT and no longer exhibited elevations in ECSIT protein found in HFD-fed Traf6^Δβ^ mice (**Figure S7B**). However, Traf6^Δβ^/ECSIT^β-het^ mice were phenotypically indistinguishable from Traf6^Δβ^ mice with regards to glucose homeostasis, insulin secretion, and insulin content when fed either RFD or HFD (**Figure S7C-J**). These data indicated that elevations in ECSIT *per se* did not elicit β-cell dysfunction following TRAF6-deficiency.

### TRAF6 mediates localization of the mitophagy machinery to mitochondria during metabolic stress

To better understand the mechanism by which TRAF6 regulates mitochondrial quality control during metabolic stress, we examined mitochondrial protein expression in β-cells by proteomics. Due to the substantial challenge of purifying sufficient mitochondrial protein from primary islets of our genetic mouse models required for proteomics profiling, we instead performed these studies in TRAF6-deficient and scramble control Min6 β-cells. Following exposure of TRAF6-deficient (or scramble control) Min6 β-cells to palmitate (or BSA, as a control), we performed biochemical fractionation to enrich for mitochondria. We then employed tandem mass tag labeling and liquid chromatography-mass spectrometry (TMT-MS) to examine mitochondrial protein expression (**Figure 5A**). We identified 764 differentially enriched proteins in the mitochondrial fraction of TRAF6-deficient β-cells compared to controls following palmitate exposure (**Figure 5B**). To identify factors associated with impairments in mitophagy following TRAF6-deficiency, we overlaid these 764 proteins with a validated list of proteins from mitophagy and autophagy pathways^58^. We identified 13 mitophagy and autophagy proteins differentially enriched in the mitochondrial fraction following TRAF6-deficiency after palmitate exposure (**Figure 5B-C**). Of these 13 proteins, 3 were significantly enriched in our TMT-MS studies with both TRAF6-deficiency and palmitate exposure, yet not by TRAF6-deficiency or palmitate treatment alone (**Figure 5B-D, Figure S8A-C**). This included the ubiquitin receptors TAX1BP1 and p62/SQSTM1, both of which participate in Parkin-dependent mitophagy and are known to interact with TRAF6^32,59,60^, and the mitophagy receptor BNIP3, which is a regulator of Parkin-independent mitophagy^61^. While all 3 proteins can be observed in other subcellular compartments, including the cytoplasm, nucleus, lysosome, and ER, these proteins are recruited to mitochondria to facilitate clearance during mitophagy^32,61,62^. We also evaluated the mitochondrial expression of other ubiquitin receptors involved in mitophagy, including CALCOCO2/NDP52, NBR1, and OPTN, as well as other mitophagy receptors, including BCL2L13, NIX/BNIP3L, and FUNDC1, and did not observe differences in mitochondrial enrichment of these proteins following TRAF6-deficiency or palmitate exposure (**Table S2**).

**Figure 5.**
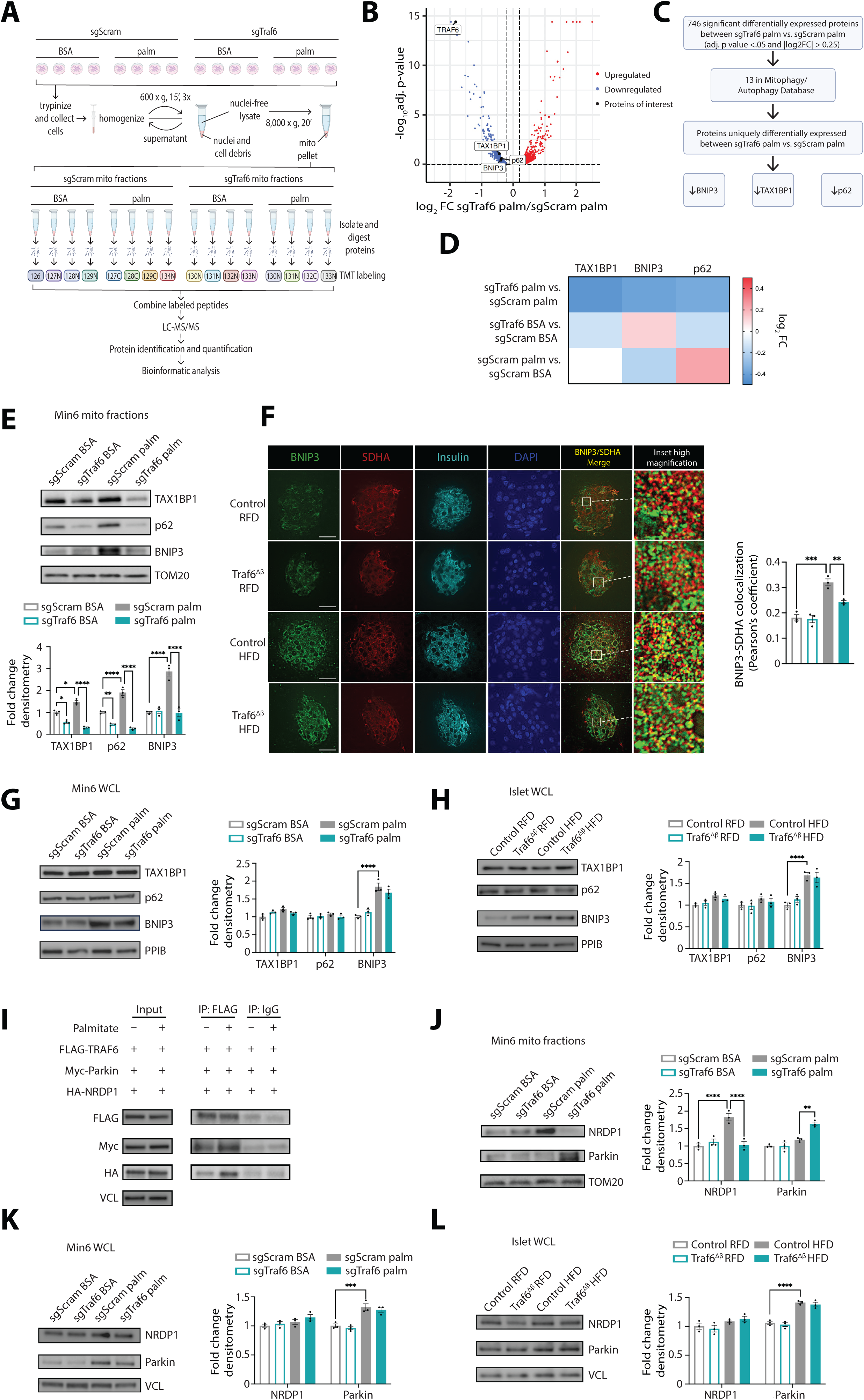
TRAF6 regulates recruitment of the mitophagy machinery following metabolic stress. **(A)** Schematic of proteomics sample preparation and experimental design. **(B)** Volcano plot of differentially expressed proteins in sgTraf6 palm vs. sgScram palm mitochondrial fractions. Differentially expressed proteins identified by adjusted *P* value < 0.05 and |log_2_FC| > 0.25. *n* = 4/group. **(C)** Workflow of protein candidate identification following proteomics analysis of mitochondrial fractions of sgTraf6 vs. sgScram Min6 cells following exposure to 0.5 mM palmitate or BSA for 48 h. **(D)** Differential protein expression heatmap of TAX1BP1, BNIP3, and p62, displayed as integrated mean log_2_FC, from proteomics studies of mitochondrial fractions of sgTraf6 vs. sgScram Min6 cells following exposure to 0.5 mM palmitate or BSA for 48 h. *n* = 4/group. **(E)** WB with densitometry of TAX1BP1, p62, and BNIP3 (relative to sgScram control) in mitochondrial fractions from sgScram and sgTraf6 Min6 cells exposed to BSA or 0.5 mM palmitate for 48 h. TOM20 served as a loading control. *n* = 3/group. \**P* < 0.05, \*\**P* < 0.01, \*\*\*\**P* < 0.0001 by 2-way ANOVA. **(F)** Representative deconvolution immunofluorescence images (left) from pancreatic sections of control or Traf6^Δβ^ mice fed RFD or HFD for 15 weeks stained for BNIP3 (green), mitochondria (SDHA; red), Insulin (cyan), and DNA (DAPI; blue). Insulin positive areas were selected and colocalization of BNIP3 and SDHA was quantified by Pearson’s correlation coefficient (right). *n* = 3/group. \*\**P* < 0.01, \*\*\**P* < 0.001 by 1-way ANOVA. Scale bar = 20 μm. **(G)** WB with densitometry of TAX1BP1, p62, and BNIP3 (relative to sgScram control) in whole cell lysates from sgScram and sgTraf6 Min6 cells exposed to BSA or 0.5 mM palmitate for 48 h. PPIB served as a loading control. *n* = 3/group. \*\*\*\**P* < 0.0001 by 2-way ANOVA. **(H)** WB with densitometry of TAX1BP1, p62, and BNIP3 (relative to Control RFD) in islets isolated from control or Traf6^Δβ^ mice fed RFD or HFD for 15 weeks. PPIB served as a loading control. *n* = 3 per group. \*\*\*\**P* < 0.0001 by 2-way ANOVA. (**I)** Representative WB of FLAG, HA, and Myc levels following anti-FLAG or anti-IgG immunoprecipitation (IP) in Min6 cells transfected with plasmids expressing FLAG-TRAF6, HA-NRDP1, and Myc-Parkin and exposed to BSA or 0.5 mM palmitate for 48 h. VCL served as a loading control. *n* = 3/group. **(J)** WB with densitometry of NRDP1 and Parkin (relative to sgScram control) in mitochondrial fractions from sgScram and sgTraf6 Min6 cells exposed to BSA or 0.5 mM palmitate for 48 h. TOM20 served as a loading control. *n* = 3/group. \*\**P* < 0.01, \*\*\*\**P* < 0.0001 by 2-way ANOVA. **(K)** WB with densitometry of NRDP1 and Parkin (relative to sgScram control) in whole cell lysates of sgScram and sgTraf6 Min6 cells exposed to BSA or 0.5 mM palmitate for 48 h. VCL serves as a loading control. *n* = 3/group. \*\*\**P* < 0.001 by 2-way ANOVA. **(L)** WB with densitometry of NRDP1 and Parkin (relative to Control RFD) in islets isolated from control or Traf6^Δβ^ mice fed RFD or HFD for 15 weeks. VCL served as a loading control. *n* = 3/ group. \*\*\*\**P* < 0.0001 by 2-way ANOVA.

We next confirmed our TMT-MS findings in mitochondrial fractions by Western blotting, observing lower levels of mitochondrial TAX1BP1, p62, and BNIP3 in TRAF6-deficient cells following palmitate exposure (**Figure 5E**). As an orthogonal approach to complement our biochemical fractionation, we isolated mitochondria utilizing magnetic sorting with an antibody specific for the mitochondrial protein TOM22^63^. Following anti-TOM22 sorting, we again confirmed reduced mitochondrial recruitment of TAX1BP1, p62, and BNIP3 in TRAF6-deficient cells following palmitate exposure (**Figure S8D**). Further, we observed lower mitochondrial BNIP3 in β-cells of HFD-fed Traf6^Δβ^ mice compared to HFD-fed control littermates, noted by lower BNIP3 colocalization with the mitochondrial marker succinate dehydrogenase A (SDHA) (**Figure 5F**).

To determine if the reductions in mitochondrial TAX1BP1, p62, and BNIP3 were related to reduced overall levels of these proteins, we next examined their expression in whole-cell extracts. Importantly, no differences in total levels of TAX1BP1, p62, and BNIP3 were observed in Min6 β-cells in the setting of TRAF6-deficiency alone or following palmitate exposure (**Figure 5G**), indicating TRAF6 regulates the mitochondrial localization rather than expression of TAX1BP1, p62, and BNIP3. Further, no differences in total expression of TAX1BP1, p62 and BNIP3 were observed in whole-cell extracts of islets from HFD-fed Traf6^Δβ^ mice compared to HFD-fed control littermates (**Figure 5H**). Total BNIP3 protein levels rose both in Min6 β-cells and islets following palmitate exposure or DIO, respectively (**Figure 5G-H**), consistent with reports of increased BNIP3 expression in response to metabolic stress^64–67^. Moreover, we observed no differences in the mRNA expression of *Tax1bp1*, *p62,* or *Bnip3* following DIO or TRAF6-deficiency (**Figure S1A and S5A**). Together, these data suggested that TRAF6 is essential for the proper recruitment of mitophagy machinery to mitochondria during metabolic stress.

Our subcellular fractionation studies, together with impaired phosphorylation of ubiquitin at Ser65 and MFN2 turnover in TRAF6-deficient β-cells during HFD (**Figure 4E-F**), led us to next explore the connections between TRAF6 and Parkin in β-cells. We observed that TRAF6 interacted with both Parkin and the E3 ligase NRDP1, which binds to Parkin to tune mitophagy and prevent abnormal mitophagic flux^68^, and that these interactions increased upon palmitate exposure

(**Figure 5I**). Mitochondrial NRDP1 localization was reduced in TRAF6-deficient Min6 cells following palmitate exposure, whereas Parkin localization to mitochondria increased (**Figures 5J, S8E**). Again, no differences in total expression of NRDP1 or Parkin were observed following TRAF6-deficiency in Min6 cells or primary islets (**Figure 5K-L**). Taken together, these results indicate that TRAF6-deficiency alters the recruitment of key regulators of both Parkin-dependent and-independent mitophagy following metabolic stress.

### Loss of Parkin restores β-cell function and glucose homeostasis following TRAF6-deficiency

Our observations of an accumulation of mitochondrial Parkin in TRAF6 deficient β-cells following metabolic stress despite impaired mitochondrial respiration and mitophagy led us to hypothesize that TRAF6 tunes mitophagic flux through regulation of Parkin. The role of Parkin is controversial in β-cells. Some studies have demonstrated that Parkin expression is essential for GSIS or recovery from palmitate-mediated damage in β-cell lines^69^. On the other hand, inappropriate or excessive engagement of Parkin-mediated mitophagy has been shown to lead to β-cell dysfunction^68,70–72^. Further, we previously demonstrated that Parkin expression in β-cells is dispensable for both glucose homeostasis and mitophagy flux at baseline and following HFD^73^, indicating that Parkin-independent mitophagy is fully capable of maintaining β-cell function following metabolic stress.

To determine whether Parkin impacts glucose homeostasis in TRAF6-deficient mice during metabolic stress, we crossed Traf6^Δβ^ mice and *Ins1*^Cre^ controls with mice bearing the *Parkin* conditional allele (*Parkin*^loxP^) to generate both dual Traf6/Parkin^Δβ^ and Parkin^Δβ^ mice, respectively. Traf6/Parkin^Δβ^ islets exhibited the expected reduction of both TRAF6 and Parkin protein levels during RFD and HFD feeding, with no respective compensatory changes in the single knockouts (**Figures 6A and S9A**). In line with previous results, Parkin was dispensable for glucose homeostasis following HFD^73^; however, deletion of Parkin in β-cells completely restored glucose homeostasis in HFD-fed TRAF6-deficient mice to the level seen in HFD-fed controls (**Figure 6B**). Additionally, GSIS was restored both in HFD-fed Traf6/Parkin^Δβ^ mice *in vivo* and in isolated islets (**Figure 6C-D**). As expected, Traf6^Δβ^, Parkin^Δβ^, Traf6/Parkin^Δβ^, and littermate controls displayed no physiological differences following RFD feeding (**Figure S9B-D**). Further, we found no differences in KCl-stimulated insulin secretion, total insulin content, or β-cell mass between Traf6^Δβ^, Parkin^Δβ^, Traf6/Parkin^Δβ^, and littermate control mice following RFD or HFD feeding (**Figure 6E-G, Figure S9E-F**). These data indicate that TRAF6 operates to keep Parkin activity in check to promote glucose homeostasis and insulin secretion during metabolic stress.

**Figure 6.**
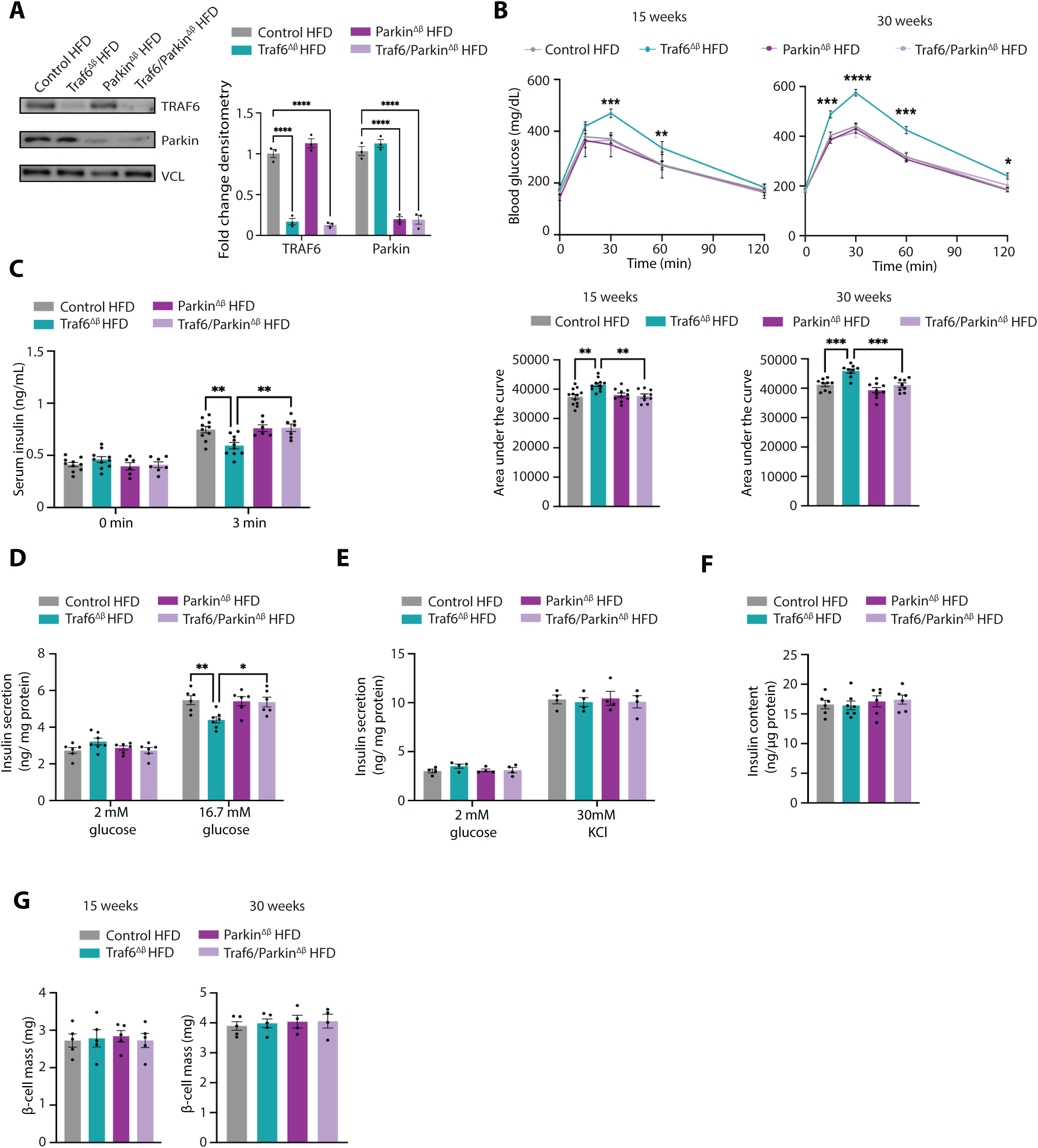
Loss of Parkin restores glucose homeostasis and β-cell function following TRAF6-deficiency. **(A)** Protein expression and densitometry quantification (relative to Control HFD) of TRAF6 and Parkin by WB in isolated islets from control, Traf6^Δβ^, Parkin^Δβ^, and Traf6/Parkin^Δβ^ mice fed HFD for 15 weeks. VCL served as a loading control. *n* = 4/group. \*\*\*\**P* < 0.0001 by 1-way ANOVA. **(B)** Blood glucose concentrations measured during IPGTT (top) and AUC (bottom) of control, Traf6^Δβ^, Parkin^Δβ^, and Traf6/Parkin^Δβ^ male mice fed HFD for 15 weeks (left) or 30 weeks (right). *n* = 9-12/group. \**P* < 0.05, \*\**P* < 0.01, \*\*\**P* < 0.001, \*\*\*\**P* < 0.0001 by 2-way ANOVA (IPGTT) or 1-way ANOVA (AUC). **(C)** Serum insulin levels measured during *in vivo* glucose-stimulated insulin release assays in male mice fed HFD for 15 weeks. *n* = 8-10/group. \*\**P* < 0.01 by 2-way ANOVA. **(D)** Glucose-stimulated insulin secretion from islets isolated from control, Traf6^Δβ^, Parkin^Δβ^, and Traf6/Parkin^Δβ^ mice fed HFD for 15 weeks. *n* = 6-7/group. \**P* < 0.05, \*\**P* < 0.01 by 2-way ANOVA. **(E)** KCl-stimulated insulin secretion from islets isolated from control, Traf6^Δβ^, Parkin^Δβ^, and Traf6/Parkin^Δβ^ mice fed HFD for 15 weeks. *n* = 4/group. **(F)** Insulin content in islets from control, Traf6^Δβ^, Parkin^Δβ^, and Traf6/Parkin^Δβ^ mice fed HFD for 15 weeks. *n* = 6-7/group. (**G)** Pancreatic β-cell mass from control, Traf6^Δβ^, Parkin^Δβ^, and Traf6/Parkin^Δβ^ mice fed HFD for 15 weeks (left) or 30 weeks (right). *n* = 4-5/group.

### Loss of Parkin rescues β-cell mitochondrial quality control following TRAF6-deficiency

We next asked whether deletion of Parkin restored the mitochondrial dysfunction and impaired mitophagy observed in HFD-fed Traf6^Δβ^ mice. Indeed, islets from HFD-fed Traf6/Parkin^Δβ^ mice exhibited a complete recovery of glucose-stimulated OCR and ATP/ADP ratio compared to islets from HFD-fed Traf6^Δβ^ mice (**Figure 7A-B**). Combined TRAF6/Parkin-deficiency restored mitochondrial mass to the same level as littermate controls, as measured via mtDNA content, citrate synthase activity, and expression of OXPHOS subunits and TOM20 protein (**Figure 7C-E**). Moreover, islets from Traf6/Parkin^Δβ^ mice also displayed rescue of both mitochondrial morphology and mitophagic flux following DIO (**Figure 7F-G**), indicating that deletion of Parkin also ameliorated Parkin-independent mitophagy in TRAF6-deficient β-cells. Indeed, mitochondrial BNIP3 localization was also improved following the loss of Parkin in HFD-fed TRAF6-deficient β-cells (**Figure 7H**). Again, no differences in mitochondrial function, morphology, or mitophagy were observed between Traf6^Δβ^, Parkin^Δβ^, Traf6/Parkin^Δβ^, and littermate control animals following RFD feeding (**Figure S9G-J**). Taken together, these results demonstrated that TRAF6 promotes ubiquitin-and receptor-mediated mitophagy in β-cells during metabolic stress by restraining Parkin activity.

**Figure 7.**
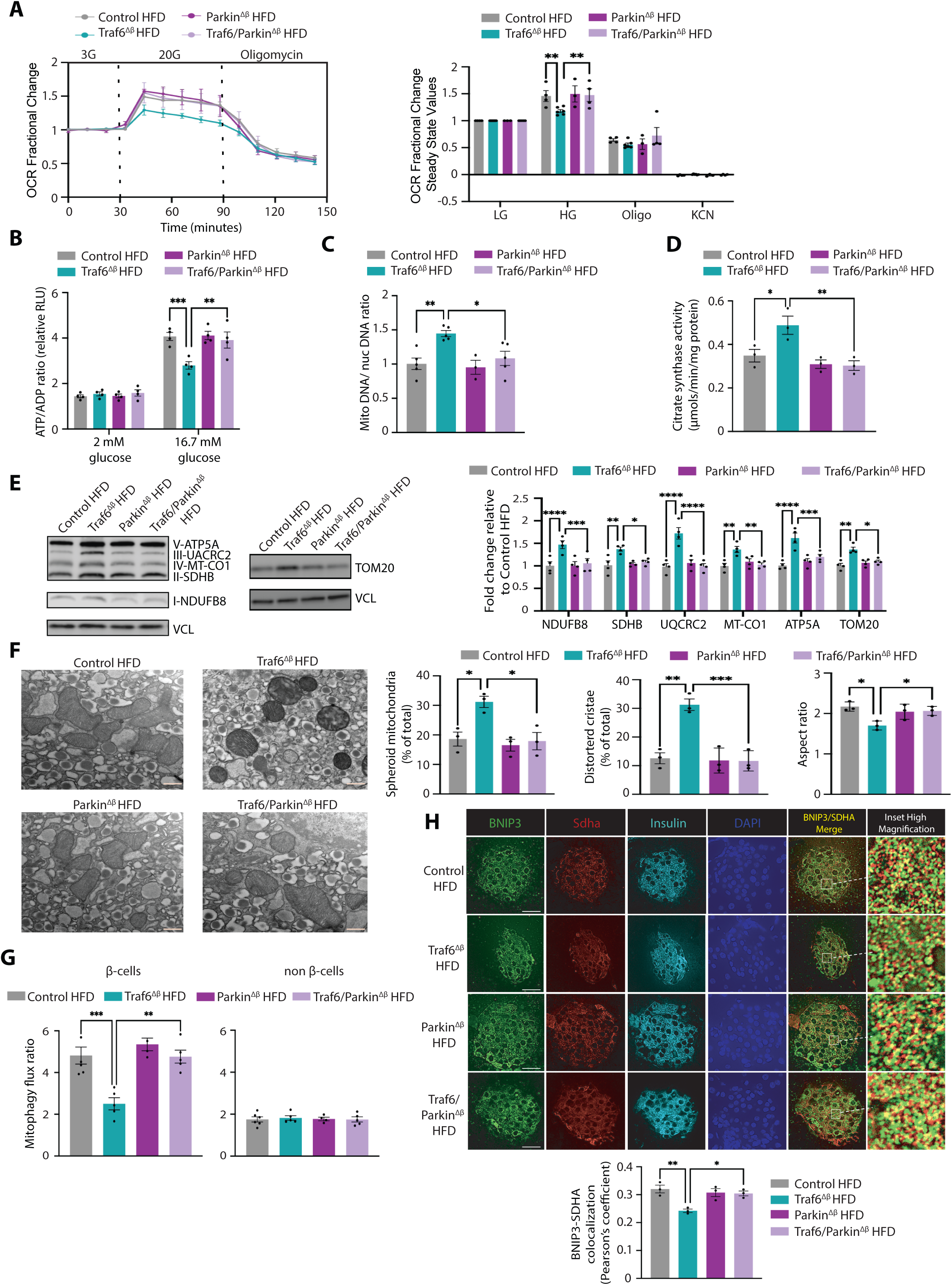
Loss of Parkin rescues β-cell mitochondrial quality control following TRAF6-deficiency. **(A)** Oxygen consumption rate (OCR) fractional change (left) and steady state quantification (right) following exposure to 3 mM glucose, 20 mM glucose, 10 μM oligomycin, and 3 mM KCN in isolated islets from control, Traf6^Δβ^, Parkin^Δβ^, and Traf6/Parkin^Δβ^ islets isolated from mice fed HFD for 15 weeks. *n* = 3-6/group. \*\**P* < 0.01 by 2-way ANOVA. **(B)** ATP/ADP ratio measured in isolated islets from control, Traf6^Δβ^, Parkin^Δβ^, and Traf6/Parkin^Δβ^ mice fed HFD for 15 weeks. *n* = 4/group. \*\**P* < 0.01, \*\*\**P* < 0.001 by 2-way ANOVA. **(C)** mtDNA to nuclear DNA ratio in isolated islets from control, Traf6^Δβ^, Parkin^Δβ^, and Traf6/Parkin^Δβ^ mice fed HFD for 15 weeks. *n* = 3-5/group. \**P* < 0.05, \*\**P* < 0.01 by 1-way ANOVA. **(D)** Citrate synthase activity in isolated islets from control, Traf6^Δβ^, Parkin^Δβ^, and Traf6/Parkin^Δβ^ mice fed HFD for 15 weeks. *n* = 3/group. \**P* < 0.05, \*\**P* < 0.01 by 1-way ANOVA. **(E)** WB for OXPHOS subunits and TOM20 with densitometry (relative to Control HFD) in islets from control, Traf6^Δβ^, Parkin^Δβ^, and Traf6/Parkin^Δβ^ mice fed HFD for 15 weeks. VCL served as a loading control. *n* = 4/group. \**P* < 0.05, \*\**P* < 0.01, \*\*\**P* < 0.001, \*\*\*\**P* < 0.0001 by 2-way ANOVA. **(F)** Representative TEM images of β-cells from control, Traf6^Δβ^, Parkin^Δβ^, and Traf6/Parkin^Δβ^ mice fed HFD for 15 weeks. with quantification of mitochondrial morphology (∼100 independent mitochondria scored/animal). *n* = 3/group. \**P* < 0.05, \*\**P* < 0.01, \*\*\**P* < 0.001 by 1-way ANOVA. Scale bars: 500 nM. **(G)** Flow cytometry quantification of mitophagy flux by MtPhagy dye approach in β-cells from control, Traf6^Δβ^, Parkin^Δβ^, and Traf6/Parkin^Δβ^ mice fed HFD for 15 weeks. *n* = 4-6/group. \*\**P* < 0.01, \*\*\**P* < 0.001 by 1-way ANOVA. (**H**) Representative deconvolution immunofluorescence images (left) from pancreatic sections of control, Traf6^Δβ^, Parkin^Δβ^, and Traf6/Parkin^Δβ^ mice HFD for 15 weeks stained for BNIP3 (green), mitochondria (SDHA; red), Insulin (cyan), and DNA (DAPI; blue). Insulin positive areas were selected and colocalization of BNIP3 and SDHA was quantified by Pearson’s correlation coefficient (right). *n* = 3/group. \**P* < 0.05, \*\**P* < 0.01 by 1-way ANOVA. Scale bars: 20 μm.

## Discussion

Here, we identified a novel link between innate immune signaling and mitochondrial quality control in pancreatic β-cells. We observed that β-cells activated innate immune signaling in response to metabolic stress. TRAF6 was dispensable for β-cell function under basal conditions, but loss of TRAF6 impaired glucose homeostasis and β-cell function following DIO. TRAF6 maintained mouse and human β-cell mitochondrial health by promoting mitophagy in the setting of metabolic stress. TRAF6-deficiency led to an aberrant increase in mitochondrial Parkin localization and impaired the recruitment of key machinery within both the Parkin-dependent and-independent mitophagy pathways during metabolic stress. Indeed, deletion of Parkin overcame both physiologic and mitophagy defects in HFD-fed Traf6^Δβ^ mice, suggesting that TRAF6 may regulate crosstalk between ubiquitin-and receptor-mediated mitophagy through Parkin to promote β-cell function (**Figure 8**). Together, our studies reveal a critical role for TRAF6 in governing adaptive β-cell responses to metabolic stressors.

**Figure 8.**
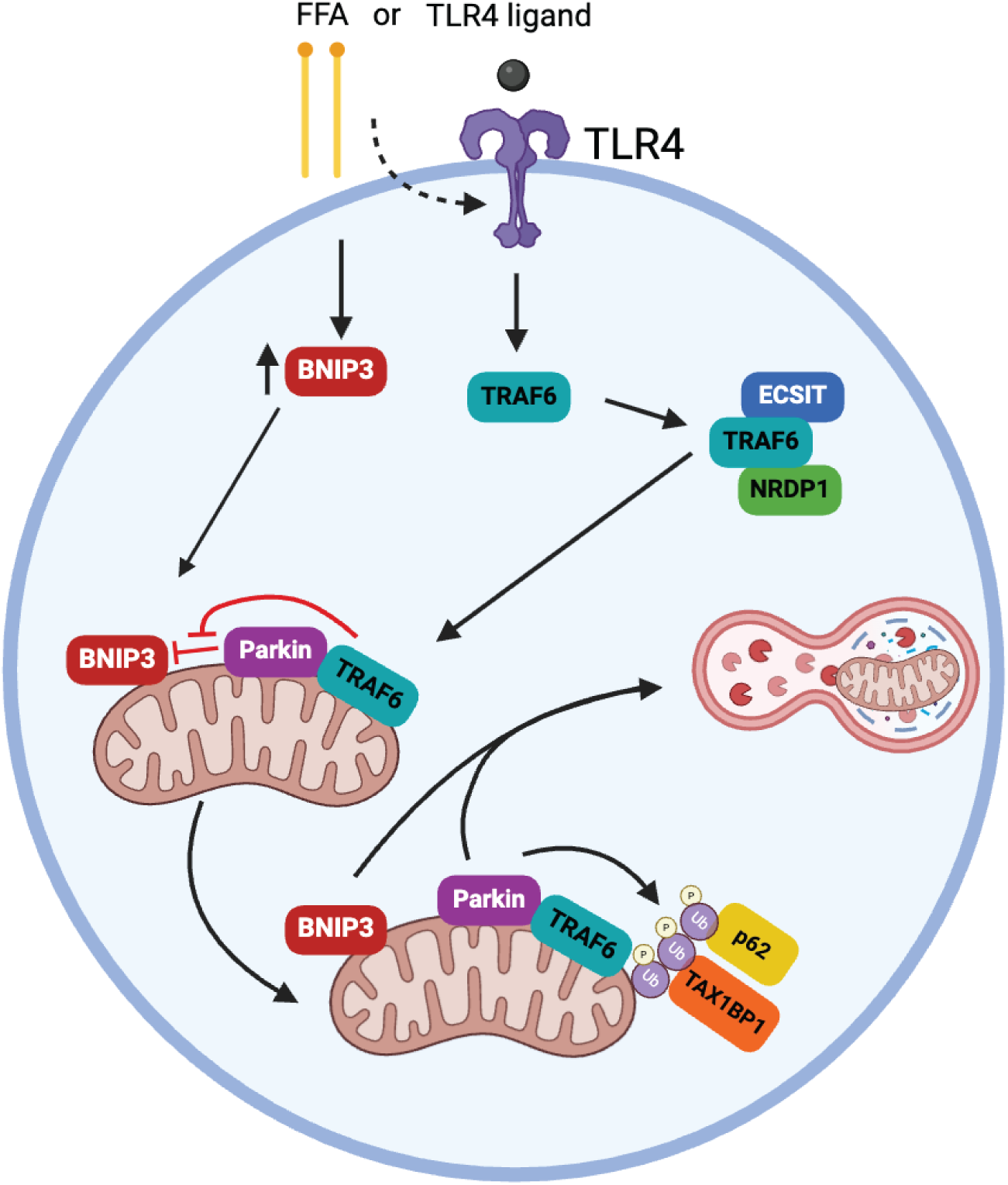
Schematic model for control of mitophagy by innate immune signaling in the adaptation to metabolic stress. Following exposure to TLR4 ligands or free fatty acids (FFA), TRAF6 interacts with ECSIT and NRDP1, and BNIP3 levels rise. TRAF6 localizes to the mitochondria where it complexes with Parkin and mediates the recruitment of ubiquitin receptors p62 and TAX1BP1. TRAF6 also restrains Parkin activity to support mitochondrial BNIP3 recruitment. Together, these events promote the induction of mitophagy to adaptively respond to metabolic stress and maintain β-cell function.

Our studies identify an unexpected intrinsic role for β-cell innate immune signaling to preserve insulin secretion in the setting of DIO or elevated free fatty acids by regulating mitophagy. An understanding for the role of innate immune signaling has expanded to non-immune cells, including studies revealing crosstalk between keratinocytes or osteoblasts with the immune system^75–79^. While we cannot eliminate a potentially similar role for TRAF6 in β-cell-immune cell crosstalk, our results suggest a β-cell-autonomous capacity for TRAF6 in directing mitochondrial quality control. Indeed, fatty acid exposure also promoted the interaction of TRAF6 with NRDP1 and ECSIT, both of which are bona fide signaling intermediates engaged during innate immune signaling with connections to mitochondrial health^55,68,80^. Our work also led us to revisit past studies of whole-body knockouts of *Trif* and *Myd88*, both upstream regulators of TRAF6 and immediately downstream of TLR4^19,20,81^, to contextualize the importance of innate immune signaling in β-cells. Indeed, *Trif-*null mice develop impaired insulin secretion, and *Myd88-*null mice are sensitized to the β-cell toxin streptozotocin^19,20^, phenotypes which bear similarity to those previously observed in models of β-cell mitophagy deficiency^72,82^. While β-cell mitophagy was not examined in studies of *Trif* and *Myd88-*null mice^19,20^, TRIF and MYD88 in macrophages interact with and activate Beclin 1, a key initiator of autophagosome formation that can also facilitate Parkin translocation to mitochondria^83–85^. However, unlike *Tlr2*/*Tlr4*-null animals^30^, Traf6^Δβ^ mice did not exhibit increases of β-cell mass following DIO. Thus, future studies will be crucial to consider the β-cell-specific relevance of numerous nodes within the innate immune signaling machinery, not only on mitophagy but also on β-cell-immune cell crosstalk to more holistically uncover their relevance to metabolic disease.

Our studies identify a novel role for TRAF6 in control of glucose homeostasis through regulation of mitophagy in β-cells that is distinct from its role in other tissues by restraining Parkin. In a model of Parkinson’s disease, Parkin promotes TRAF6 degradation, and Parkin-deficiency leads to apoptosis due to stabilization of TRAF6^87^. In neuroblastoma and HEK293 cells, TRAF6 stabilizes PINK1 on damaged mitochondria to recruit Parkin for activation of mitophagy^88^. Further, TRAF6 was previously shown to interact with p62 to regulate macroautophagy in the immune system and cancer cells^60,89^, and with TAX1BP1 to regulate NF-κB transcriptional activation in immune cells^59^. Despite a similar series of partners, our work suggests TRAF6 and these partners have distinct functions in β-cells during metabolic stress. Indeed, we observed that TRAF6-deficiency led to impaired mitophagy flux, without overt defects in macroautophagy. Despite a reduction in phospho-Ser65-ubiquitin in TRAF6-deficient β-cells, which could be suggestive of impaired PINK1 function, we observed an increase, and not a decrease, of mitochondrial Parkin following TRAF6-deficiency. We also observed TRAF6 was crucial for mitochondrial recruitment of p62 and TAX1BP1, both ubiquitin receptors in the Parkin-dependent mitophagy pathway, following metabolic stress. While Parkin-deficiency itself did not yield metabolic dysfunction or loss of β-cell mass or regulate TRAF6 expression, co-deletion of Parkin with TRAF6 reversed the glucose intolerance observed in HFD-fed Traf6^Δβ^ mice. These results suggest that the absence of TRAF6 removes a brake on Parkin activity in stressed β-cells. Tight control of Parkin activity has been suggested to be important for β-cell function^71,72^; however, the mechanisms underlying restraint of Parkin action had been previously unclear. Together, these results suggest that TRAF6 plays a critical role in allowing β-cells to adapt to metabolic stress by integrating innate immune signals and directing mitophagic flux through a complex network of partners. Additional studies will be valuable to clarify the intricacies of both tissue-and context-specific roles of TRAF6 and its numerous partners in directing mitochondrial quality control following the engagement of innate immune signaling.

Direct mechanisms underlying the crosstalk between ubiquitin-and receptor-mediated mitophagy are emerging yet remain unclear. Our work reveals the importance of both mitophagy pathways in β-cells, as well as novel crosstalk between Parkin and receptor-mediated mitophagy, which frequently is characterized as Parkin-independent. PINK1-and Parkin-deficiency are each dispensable for mitophagy in β-cells^73,90^, supporting the ability of PINK1/Parkin-independent pathways to maintain mitochondrial quality control. However, our work shows that the dispensable nature of Parkin does not render its function unimportant in β-cells and that Parkin activity must be finely tuned to balance both mitophagy pathways for optimal mitochondrial adaptation to metabolic stress. Studies of receptor-mediated mitophagy in β-cells are more limited but have recently begun to surface. Induction of BNIP3 expression by HIF1⍺ was recently shown to be vital for increases in β-cell mitophagy following DIO^64^. We similarly observe an increase in BNIP3 expression following DIO but demonstrate that TRAF6 regulates the mitochondrial recruitment rather than the expression of BNIP3, suggesting that HIF1⍺ and TRAF6 may play distinct roles in BNIP3-dependent mitophagy following DIO. While a previous study demonstrated BNIP3-dependent mitophagy is reduced following loss of Parkin in cardiomyocytes^91^, it is unclear how unchecked Parkin activity and impaired Parkin-dependent mitophagy following TRAF6 deficiency regulates crosstalk with the BNIP3 pathway. Future studies will be important to determine if the restoration of mitochondrial BNIP3 is required for the recovery of mitophagy following combined TRAF6/Parkin deficiency. While we did not observe alterations in the recruitment of other mitophagy receptors following TRAF6 deficiency, including NIX, FUNDC1, or BCL2L13^92–94^, it is also possible that defects in mitophagy (and rescue following concomitant Parkin-deficiency) could be mediated by the activity of these receptors, whose roles in β-cell mitophagy are unknown. Together, our work provides novel insights into the cross-regulation of these two distinct pathways underlying mitochondrial quality control by TRAF6 via restriction of Parkin.

T2D is a complex, multi-organ disease driven by β-cell dysfunction and impaired responses to immunometabolic stress^27,95^. While lipotoxicity is a precipitating factor for β-cell damage in T2D^96^, our studies suggest that innate immune signaling forms a pivotal protective response to equip β-cells with the means to compensate for metabolic stress, rather than as a purveyor of β-cell demise. The protective effects of innate immune signaling appear to be due to a non-canonical upregulation of mitophagy as opposed to classical transcriptional effects, but this remains to be explored in T2D. Further, it will be interesting to determine if recently reported defects in β-cell mitophagy in T2D are due to impairments in innate immune signaling^36^. Moreover, the effects of enhancement of β-cell innate immune signaling or TRAF6-mediated mitophagy could be of interest for therapeutic benefit in T2D. Together, our studies suggest that activation of innate immune signaling in β-cells is a crucial adaptive response to obesity that promotes mitophagy to preserve β-cell function.

## Materials and Methods

### Genetically modified mouse lines

All mice were maintained in accordance with the University of Michigan’s Institutional Animal Care and Use Committee under specific pathogen-free conditions. Up to 5 mice were housed per cage and on a 12 h light-dark cycle. *Traf6^loxP/loxP^* mice possessing loxP sites flanking exon 7 of the *Traf6* gene were obtained from the Jackson Laboratories (Traf6 Strain 030849)^97^. *Parkin*^fl/fl^ mice possessing loxP sites flanking exon 7 of the *Parkin* gene were generously provided by Dr. Ted Dawson (Johns Hopkins University) and Lexicon Genetics^74^. *Ecsit*^fl/fl^ mice containing a loxP site flanking exon 3 were previously described^55^. To generate β-cell-specific deletion, floxed models were intercrossed with *Ins1*-Cre knockin mice obtained from Jackson laboratories (JAX Stock No. 026801)^98^. *Ins1*-Cre–alone and respective floxed-only controls for each study (*Traf6^loxP/loxP^*, *Parkin^fl/fl^*, *Traf6^loxP/loxP^/Parkin^fl/fl^*or *ECSIT^fl/fl^* mice) were phenotypically indistinguishable from each other and combined as controls. *Ins1*-Cre–alone were also phenotypically indistinguishable from wild-type C57BL/6N controls, consistent with previous reports from our group and others^36,73,82,98,99^. mt-Keima mice were a gift from Dr. Toren Finkel (University of Pittsburgh)^50^. All mouse models were maintained on a 100% C57BL/6N background. All studies and endpoints were completed using both male and female mice.

### Human islet samples

All human samples were procured from de-identified donors from Prodo Laboratories or the Alberta Diabetes Institute IsletCore. Studies were approved by the University of Michigan Institutional Review Board. Human primary islets were cultured at 37 **°**C with 5% CO_2_ in PIM(S) media (Prodo Laboratories) supplemented with 10% FBS (Gemini Bio), 100 U/mL penicillin/streptomycin (Gibco), 100 U/mL antibiotic/antimycotic (Gibco), and 1mM PIM(G) (Prodo Laboratories). Islets were used from male and female donors, and donor information is provided in **Table S3**. Drug treatments used for human islet cultures included 1 μM C25-120 (TRAF6-UBC13 inhibitor, 24 h, Med Chem Express^100^), 250 nM valinomycin (3 h, MilliporeSigma), or glucolipotoxicity (25 mM glucose + 0.5 mM palmitate (conjugated to BSA as previously described^73^), 48h), as well as respective vehicle controls (DMSO or BSA).

### High-fat diet feeding

Control, Traf6^Δβ^, ECSIT^β-het^, Traf6^Δβ^/ECSIT^β-het^, Parkin^Δβ^, and Traf6/Parkin^Δβ^ mice were randomized onto a 10% regular-fat diet or 60% high-fat diet (Research Diets Inc) at weaning. Access to food was *ad libitum* for up to 30 weeks. Mice were then subjected to *in vivo* metabolic analyses (detailed in following sections), and pancreatic tissue was harvested at the end of the study for immunohistochemistry, *ex vivo* islet studies, metabolic phenotyping or sequencing studies. Physiologic data in male and female mice cohorts were displayed separately to evaluate for potential sexual dimorphic metabolic phenotypes following diet-induced obesity.

### Mouse primary islet isolation and culture

Mouse primary islets were isolated by perfusing pancreata with a 1 mg/mL solution of Collagenase P (Millipore Sigma) in 1 X HBSS (Sigma) into the pancreatic duct. Following excision, pancreata were incubated at 37 °C for 13 min, and Collagenase P was deactivated by addition of Quench Buffer (1X HBSS + 10% adult bovine serum (Gemini Bio)). Pancreata were dissociated mechanically by vigorous shaking for 30 sec, and the resulting cell suspension was passed through a 70 μM cell strainer (Fisher Scientific). Cells were centrifuged at 1000 rpm for 2 min, then the pellet was resuspended in 20 mL Quench buffer and gently vortexed to thoroughly mix. Cells were again centrifuged at 1000 rpm, 1 min. This wash step was repeated once more. Following washes, the cell pellet was resuspended in 5 mL Histopaque (Millipore-Sigma) with gentle vortexing. An additional 5 mL Histopaque was layered on the cell suspension, and 10 mL Quench buffer was gently layered on top. The cells were spun at 800 x *g* for 30 min at 10 °C, with the brake off. The entire Histopaque gradient was pipetted off and passed through an inverted 70 μM filter to trap the islets cells. Islets were washed twice with 10 mL of Quench buffer and once with 10 mL of islet media (RPMI-1640 supplemented with 100 U/mL penicillin/streptomycin, 10% FBS, 1 mM HEPES, 2 mM L-Glutamine, 100 U/mL antibiotic/antimycotic and 10 mM sodium pyruvate). The filter was inverted into a sterile petri dish and cells were washed into the dish with 4.5 mL complete islet media. Islets were left to recover overnight, and treatments began the next day. Mouse islets were treated with the following treatments: 100 nM, 10 nM, or 1 μM of C25-120 (TRAF6-UBC13 inhibitor, 24 h, Med Chem Express); Valinomycin (250 nM, 3 h, Sigma Aldrich); Lipopolysaccharides from *Escherichia coli* O55:B5 (LPS) (5 ng/mL, 1 or 2 h, Sigma); pro-inflammatory cytokine cocktail (75 U/mL IL-1β, 750 U/mL TNF-α, and 750 U/mL IFN-γ, 6 h, Peprotech); palmitate (0.5 mM palmitate, 24 or 48 h, Sigma Aldrich); or vehicle controls. Islets were isolated from both male and female mice.

### Cell culture, cell line generation, treatments, and transfections

Min6 cells (a gift from Dr. Doris Stoffers, University of Pennsylvania) were cultured as previously described^73^. *Traf6*-deficient Min6 cells were generated using the LentiCRISPRV2 one-vector system as described^101^. *Traf6* sgRNA sequences included (5’-GGAGATCCAGGGCTACGATG-3’, targeting exon 3) and (5’-ATTTGGGCACTTTACCGTCA-3’ targeting exon4) and were cloned into LentiCRISPRV2 as described^102^. LentiCRISPRV2 was transfected into 293 T cells along with the lentiviral packaging plasmids RRE, VSV-G, and REV (courtesy of Dr. Ling Qi, University of Virginia) using Lipofectamine 2000 (Invitrogen). Viral-containing media were collected on days 3 and 4 post-transfection, filtered in a 0.45-µm filter, and stored at −80**°** C until use. Min6 cells were transduced by culturing with viral media (DMEM (Gibco) supplemented with 10% volume fetal bovine serum (Gemini Bio Products), 50 units/mL penicillin streptomycin (Thermo Fisher Scientific), and 10 mM HEPES (Gibco)), pH 7–7.4 mixed 1:1 with normal Min6 media (as previously described^101^), supplemented with polybrene (5 µg/mL, Sigma). Fresh viral stocks were added daily for two days, followed by two rounds of selection using puromycin (2 µg/mL, Cayman Chemical) for 5 days. Min6 cells transduced with *Traf6* sgRNA LentiCRISPR V2 (or non-targeting control sgRNA, courtesy of Dr. Ling Qi, University of Virginia) were screened via western blot.

### Luciferase assay

Either sgScramble or sgTraf6 Min6 cells were split with 150,000 cells per well in 24-well black well clear bottom clear bottom TC plates (Revvity). The next day, cells were transfected with 500 μg of 15 pathway reporter vectors or empty vector control (pLenti background with both GFP and firefly luciferase promoters) using Lipofectamine 3000^37^. Pathway reporter vectors are listed in **Table S1**^37^. After 24 hours, cells were treated with LPS (5 ng/mL, 2 h, Sigma), palmitate (0.5 mM, 48 h, Sigma Aldrich), or vehicle controls. Following treatments, cells were placed in phenol-free RPMI media (Gibco) and GFP intensity was measured using a fluorescence plate reader by measuring excitation wavelength at 485 nm and emission wavelength at 528 nm. After, cells were scraped and lysed using 1X lysis buffer (Promega Catalog# E4030). To measure relative light units, the Veritas luminometer with injector was used. 20 μL of cell lysate was loaded into white-bottom 96 well plates, and 100 μL of luciferase substrate (Promega) was injected per sample. Luminescence was measured using a 2 second delay and 8 second read time. Data were presented as relative light units (RLU) normalized to GFP intensity.

### RNA-seq on primary mouse islets

Sequencing was performed by the Advanced Genomics Core at University of Michigan Medical School. Total RNA was isolated from isolated islets and DNase treated using commercially available kits (Omega Biotek and Ambion). Libraries were constructed and subsequently subjected to 151 bp paired-end cycles on the NovaSeq-6000 platform (Illumina). FastQC (v0.11.8) was used to ensure the quality of data. Reads were mapped to the reference genome GRCm38 (ENSEMBL), using STAR (v2.6.1b) and assigned count estimates to genes with RSEM (v1.3.1). Alignment options followed ENCODE standards for RNA-seq. FastQC was used in an additional post-alignment step to ensure that only high-quality data were used for expression quantitation and differential expression. Data were pre-filtered to remove genes with 0 counts in all samples. Differential gene expression analysis was performed using DESeq2, using a negative binomial generalized linear model (thresholds: linear fold change > 1.5 or <-1.5, Benjamini-Hochberg FDR (P_adj_) <0.05). Plots were generated using variations of DESeq2 plotting functions and other packages with Genialis.

### Intraperitoneal glucose tolerance tests (IPGTT)

Mice were fasted for 6 h. Fasting blood glucose measurements were taken by tail nick (Bayer Contour) before an IP injection of 2 g/kg glucose was administered for all mice except males fed HFD, who received 1 g/kg. Blood glucose measurements were then taken at 15, 30, 60 and 120 mins. Following the test, mice were returned to housing cages with *ad libitum* access to food.

### *In vivo* glucose stimulated insulin release

Mice were fasted for 6 h. Fasting blood glucose was measured after tail nick with a glucometer (Bayer Contour) and a 20 μL blood sample was collected using capillary tubes (Fisher Scientific) and stored on ice. Mice were injected with 1.5 mg/kg glucose and blood glucose and blood samples were taken after 3 min. Blood samples were ejected from the capillary tubes into 1.5 mL tubes, spun at 15,000 x *g*, 4 °C for 10 min, and serum was aliquoted to new 1.5 mL tubes. Serum insulin levels were measured by ELISA (Alpco). Following the test, mice were returned to housing cages with *ad libitum* access to food.

### Intraperitoneal insulin tolerance tests (ITT)

Mice were fasted for 6 h. Fasting blood glucose was measured after tail nick with a glucometer (Bayer Contour). Mice were injected with 0.8 U/kg insulin (Humulin R, Eli Lilly) and blood glucose measured at 0, 15 and 30 min. Following the test, mice were returned to housing cages with *ad libitum* access to food.

### Western blot

Mouse pancreatic islets were lysed with RIPA buffer containing protease and phosphatase inhibitors (Calbiochem), and insoluble material was removed by centrifugation. Equal amounts of proteins were resolved on 4%–20% gradient Tris-glycine gels (Bio-Rad) and transferred to nitrocellulose membranes (Bio-Rad). Membranes were then blocked in 5% milk for 1 h and immunoblotting was performed using BNIP3 (1:1000; Abcam, Catalog#ab109362), Cyclophilin B (PPIB) (1:5000; ThermoFisher, Catalog#PA1-027A), ECSIT (1:1000; Abcam, Catalog#ab21288), FLAG clone M2 (1:1000; Sigma, Catalog# F104), HA (1:1000, Santa Cruz, Catalog#sc-805), LC3 (1:1000; Sigma, Catalog#L-8919), MFN2 (1:1000; Abcam, Catalog#ab56889), myc-HRP (1:1000; Roche, Catalog#a790-4628), NDUFA9 (1:1000; Abcam,Catalog#ab14713), NRDP1 (1:1000; Santa Cruz, Catalog#sc-365622), Total OXPHOS (1:1000; Abcam, Catalog#ab110413), p62 (1:1000; Enzo, Catalog#BML-PW9860), Parkin (1:500, Cell Signaling, Catalog#4211), Phospho-(Ser65)-ubiquitin (1:1000; Cell Signaling, Catalog#62802), TAX1BP1 (1:1000; Proteintech, Catalog#14424-1-AP), TOM20 (1:1000; Cell Signaling Technology, Catalog#42406), TRAF6 (1:1000; Abcam, Catalog#ab40675), Vinculin (VCL) (1:5000; Millipore, Catalog#CP74) and species-specific HRP-conjugated secondary antibodies (Vector Laboratories). Antibodies are listed in **Table S4**.

### Quantitative PCR and mtDNA assessment in primary mouse islets

DNA and RNA samples were isolated from mouse islets with the Blood/Tissue DNeasy kit (Qiagen) and MicroElute total RNA kit (Omega Biotek), respectively. DNase-treated RNA (Ambion DNA free kit) was reverse transcribed using High-Capacity cDNA Reverse Transcription Kit (Thermo Fisher Scientific) as previously described^103^. Quantitative PCR (qPCR) was performed with SYBR-based detection (Universal SYBR Green Supermix, Bio-Rad). Relative mtDNA content was quantified by qPCR using mtDNA specific primers *Mt9* and *Mt11*, and nuclear DNA specific primers *Ndufv1* Forward and Reverse. For gene expression analysis of cDNA, primers included *Traf6* and *Hprt*. qRT-PCR primer sequences are listed in **Table S5**.

### ATP/ADP ratio assessments in primary mouse islets

Isolated islets from mice were cultured overnight at 37 C. On the day of the experiment, 75 islets per group (low and high glucose for each genotype) were picked and placed in 2mL 1x KRB (1mM CaCl_2_, 4.7 mM KCl, 5 mM KH_2_PO_4_, 1mM MgSO_4_, 136 mM NaCl, 20 mM HEPES pH 7.4) for 30 min at 37 C. For the high glucose group, 200 μL of KRB solution is removed and replaced with 200 μL of 200 mM glucose solution to bring glucose concentration to 20 mM. Islets were incubated for 5 min a 37 C, then picked and collected in a microcentrifuge tube. Islets were then washed twice with 1x PBS and then dispersed into single cells using 500 μL 0.05% trypsin and incubation at 37 °C for 3 min. This was followed by gentle pipetting up and down approximately 10 times and neutralization of the trypsin with mouse islet media (RPMI media supplemented with glutamine, antimycotic-antibiotic, sodium pyruvate, and 10% BSA). Single cells were sedimented by centrifugation at room temperature, 2000 rpm, 1 min, and the pelleted cells were washed twice with 1X PBS. The cells were resuspended in 10 μL 1x PBS. Single cells were pipetted into a 96-well white well clear bottom clear bottom plate. Single cell suspensions were used to determine the ATP/ADP ratio using the ATP/ADP ratio assay kit (Sigma-Aldrich) per manufacturer’s instructions. Luminescence was measured using a 5 second integration time.

### Citrate synthase activity in primary mouse islets

Mouse islets were lysed with RIPA buffer containing protease and phosphatase inhibitors (Calbiochem), and insoluble material was removed by centrifugation. Lysates were used to determine citrate synthase activity with the MitoCheck Citrate Synthase Activity Assay Kit (Cayman Chemical) per manufacturer’s instructions.

### Mitophagy assessment in primary mouse and human islets using MtPhagy dye

Isolated islets from mice were cultured overnight at 37 °C. On the day of the experiment, 100 islets were picked per condition and treated with either vehicle (DMSO) or 250 nM valinomycin (Sigma) for 3 h. Islets were then washed twice with 1x PBS and dispersed into single cells using 500 μL 0.05% trypsin and incubation at 37 °C for 3 min. This was followed by gentle pipetting up and down approximately 10 times and neutralization of the trypsin with mouse islet FACS media (phenol-free RPMI media supplemented with glutamine, antimycotic-antibiotic, sodium pyruvate, and 10% BSA). Single cells were sedimented by centrifugation at room temperature, 2000 rpm, 1 min, and the pelleted cells were washed twice with 1X PBS. The cells were resuspended in 500 μL mouse islet FACS buffer. Cells were incubated with 500 nM Fluozin-3AM (Thermofisher), 100 nM MtPhagy Dye (Dojindo), and 100 nM TMRE (Anaspec) for 30 min at 37 C to label insulin granule positive β cells, cells undergoing mitophagy, and mitochondrial membrane potential, respectively. Cells were centrifuged at 1400 rpm at room temperature for 3 min and resuspended in 500 μL islet FACS buffer, and 0.2 μg/mL DAPI was added. Samples were analyzed on an Attune Nxt cytometer (Thermofisher). Single cells were gated using forward scatter and side scatter (FSC and SSC, respectively) plots, DAPI staining was used to exclude dead cells, and Fluozin-3AM was used to identify β-cells. MtPhagy measurements in β-cells were made using 488 nm excitation laser with a 710 nm emission filter and analyzed using FlowJo (Tree Star Inc.). A total of 5,000 β-cells was quantified from each independent islet preparation. Mitophagy flux was calculated using a ratio of MtPhagy^high^/TMRE^low^ cells exposed to valinomycin to MtPhagy^high^/TMRE^low^ cells not exposed to valinomycin. The gating strategy was described previously^49^. Human islets from non-diabetic donors were cultured for 48 h after shipment. Upon receiving the islets, islets were treated with either control (BSA, 48 h) or glucolipotoxicity (GLT; 25 mM glucose + 0.5 mM palmitate, 48 h). After 24 h, islets were treated with either vehicle (DMSO, 24 h) or 1 μM C25-120 (TRAF6-UBC13 inhibitor, 24 h, MedChem Express) in addition to the control or GLT. Preparation for flow cytometry was described as above.

### Mitophagy assessments in primary mouse islets using mt-Keima

Islets were isolated from mt-Keima mice fed either 15-week RFD or HFD and incubated overnight at 37 °C. Islets were treated with either vehicle (DMSO, 24 h) or 1 μM C25-120 (TRAF6-UBC13 inhibitor, 24 h, MedChem Express). On the day of the experiment, 100 islets were picked per condition and treated with either Vehicle (DMSO) or 250 nM Valinomycin (Sigma) for 3 h. Islets were then washed twice with 1x PBS and then dispersed into single cells using 500 μL 0.05% trypsin and incubation at 37 °C for 3 min. This was followed by gentle pipetting up and down approximately 10 times and neutralization of the trypsin with mouse islet FACS buffer. Single cells were stained with DAPI (Thermo Fisher Scientific) and resuspended in 500 μL of islet FACS media and 0.2 μg/mL DAPI was added. Samples were analyzed on an Attune Nxt cytometer (Thermofisher). Single cells were gated using forward scatter and side scatter (FSC and SSC, respectively) plots, and DAPI staining was used to exclude dead cells. Dual-excitation ratiometric mt-Keima measurements were made using 488 nm (neutral) and 561 nm (acidic) excitation lasers with 610 nm emission filter^50,104^. Ratios of acidic to neutral cell populations were then calculated using FlowJo (Tree Star Inc.). A total of 5,000 β-cells were quantified from each independent islet preparation. Mitophagy flux was calculated using a ratio of the acidic/neutral cells exposed to valinomycin to ratio of acidic/neutral cells not exposed to valinomycin. The gating strategy was described previously^49^.

### β-cell mass analysis

Whole mouse pancreas was excised, weighed and fixed in 4% paraformaldehyde for 16 h at 4 °C. Samples were stored in 70% ethanol, 4 °C before being embedded in paraffin and sectioned. 3 independent depths of sections, at least 50 μM apart, were dewaxed, and rehydrated and antigen retrieval was carried out using 10 mM sodium citrate (pH 6.0) in a microwave for 10 min. Sections were washed twice with 1X PBS, blocked for 1 h at room temperature with 5% donkey serum in PBT (1X PBS, 0.1% Triton X-100, 1% BSA). Sections were then incubated in primary insulin antibody (1:500; Invitrogen, Catalog#701265). Sections were then washed twice with PBS and incubated for 2 h at room temperature with a species-specific Cy2 secondary antibody. Nuclear labelling was performed using DAPI (Molecular Probes). Sections were scanned using an Olympus IX81 microscope (Olympus) at 10X magnification, with image stitching for quantification. β-cell mass quantification (estimated as total insulin positive area/total pancreatic area multiplied by pancreatic weight) was performed on stitched images of complete pancreatic sections from 3 independent regions.

### Colocalization studies

Whole mouse pancreas was excised and fixed in 4% paraformaldehyde for 16 h at 4 °C. Samples were stored in 70% ethanol, 4° C before being embedded in paraffin and sectioned. Slides were dewaxed and rehydrated, and antigen retrieval was carried out using 10 mM sodium citrate (pH 6.0) in a microwave for 10 min. Sections were washed twice with 1X PBS, blocked for 1 h at room temperature with 5% donkey serum in PBT (1X PBS, 0.1% Triton X-100, 1% BSA). Sections were then incubated in the following primary antisera overnight at 4C in PBT: BNIP3 (1:100; Abcam, Catalog#ab109362), insulin (1:1000; Fitzgerald, Catalog#20-IP35), and SDHA (1:100; Abcam, Catalog#ab14715). Sections were then washed twice with PBS and incubated for 2 h at room temperature with species specific Cy2, Cy3, and Cy5 conjugated secondary antibodies (Jackson Immunoresearch). Nuclear labelling was performed using DAPI (Molecular Probes). *Z*-stack images were captured with an IX81 microscope (Olympus) at 100x magnification using an ORCA Flash4 CMOS digital camera (Hamamatsu) and subjected to deconvolution (CellSens; Olympus). The BIOP-implemented interpretation of the FIJI-plugin JaCOP was used to derive Pearson’s coefficients as a measure of colocalization, selecting only insulin positive areas as the region of interest^105,106^.

### Islet respirometry

Islet respirometry was performed using the BaroFuse instrument (EnTox Sciences, Inc) as previously described^107^. Setup of the BaroFuse perifusion instrument involved 4 steps: 1. The system was preheated by filling the Media Reservoirs with pre-equilibrated Media (1X KRBH with 3 mM glucose); 2. The Tissue Chamber Assemblies were inserted into the Perifusion Module and secured in place on top of the Media Reservoir to create a gas tight seal; 3. The headspace of the Media Reservoirs were purged with the desired gas composition (typically 5% CO_2_/21% O_2_ balance N_2_) to allow the gas in the headspace to equilibrate with the perifusate in the reservoir; and 4. The purge port was closed to allow the flow to fill up the Tissue Chambers. When the Tissue Chambers had almost filled, islets were loaded into 6 of the 8 chambers and 2 were left empty to be used as an inflow reference. 100 islets were transferred into the Tissue Chamber (ID = 1.5 mm) with a P-200 pipet. Once islets were loaded, the magnetic stirrers, O2 detector and flow rate monitor were started, and the system and the islets were allowed to equilibrate to establish a stable baseline for 90-120 minutes. Subsequently, test compounds (final concentrations of 20 mM glucose, 10 μM oligomycin, and 3 mM KCN) were injected at precise times. Data are presented as fractional change from baseline (at 3 mM glucose containing media) oxygen concentration rates (OCR) per sample relative to minimum OCR per sample determined following exposure to 3 mM KCN for baseline normalization using the BaroFuse Data Processing Package software (Entox Sciences, Inc.).

### Transmission electron microscopy

Mouse islets were fixed in 2% paraformaldehyde and 2.5% glutaraldehyde in 0.1 M sodium cacodylate (pH 7.4) for 5 minutes at room temperature and then stored at 4**°** C for further processing. Following agarose embedding, islets were then subjected to osmification in 1.5% K_4_Fe(CN)_6_ + 2% OsO_4_ in 0.1 cacodylate buffer for 1 h, dehydrated by serial diluted ethanol solutions (30, 50, 70, 80, 90, 95 and 100%) and embedded in Spurr’s resin by polymerization at 60**°** C for 24 h. Polymerized resins were then sectioned at 90 nm thickness using a Leica EM UC7 ultramicrotome and imaged at 70 kV by using a JEOL 1400 transmission electron microscopy equipped with an AMT CMOS imaging system. Mitochondrial morphology was analyzed and quantified by Fiji, as previously described^99^.

### ROS assessments

ROS levels were measured via flow cytometry in isolated islets by a cell-based fluorogenic assay kit (ROS/Superoxide detection assay kit, Abcam) as per the manufacturer’s protocol as previously described^73^.

### Plasmids

Constructs for co-immunoprecipitation overexpression studies included pCMV5-FLAG-TRAF6^108^, pCMV3-C-HA-ECSIT (Sino Biological, Catalog#HG14497-CY), pRK5-Myc-Parkin^109^, and pcDNA3.1 3x-HA-NRDP1^110^. FLAG-TRAF6-wt was a gift from John Kyriakis (Addgene plasmid #21624) and pRK5-Myc-Parkin was a gift from Ted Dawson (Addgene plasmid #17612). 2 μg of constructs were transfected into Min6 cells using XtremeGene transfection reagent (Millipore Sigma). For luciferase overexpression studies, pLminP_Luc2P_RE1 (Addgene plasmid #90335), pLminP_Luc2P_RE4 (Addgene plasmid #90338), pLminP_Luc2P_RE5 (Addgene plasmid #90339), pLminP_Luc2P_RE10 (Addgene plasmid #90344), pLminP_Luc2P_RE21 (Addgene plasmid #90363), pLminP_Luc2P_RE23 (Addgene plasmid #90365), pLminP_Luc2P_RE26 (Addgene plasmid #90368), pLminP_Luc2P_RE27 (Addgene plasmid #90369), pLminP_Luc2P_RE28 (Addgene plasmid #90370), pLminP_Luc2P_RE34 (Addgene plasmid #90376), pLminP_Luc2P_RE38 (Addgene plasmid #90378), pLminP_Luc2P_RE47 (Addgene plasmid #90385), pLminP_Luc2P_RE52 (Addgene plasmid #90397), and pLminP_Luc2P_RE57 (Addgene plasmid #90402) were gifts from Ramnik Xavier^37^.

### Immunoprecipitation studies

Min6 cells were transfected with FLAG-TRAF6, HA-ECSIT, HA-NRDP1, and Myc-Parkin constructs using XtremeGene transfection reagent (Millipore Sigma). The following day, cells were treated with either BSA control or 0.5 mM palmitate for 48 h. Cells were trypsinized and collected, then washed twice with 1x PBS. Cells were then lysed in a protein lysis buffer consisting of 150 mM NaCl, 1% IGEPAL CA-630, and 50 mM Tris, pH 8.0, supplemented with protease and phosphatase inhibitors (Millipore), followed by shearing via passage through a 21-gauge needle while on ice. Lysates were clarified by centrifugation, then pre-cleared with protein G agarose beads (Thermo Fisher Scientific, Catalog#20399). 500 μg of protein lysates were incubated with FLAG M2 Affinity Gel (Sigma Aldrich, Catalog#A2220) overnight at 4° C. Beads were washed 5 times in lysis buffer, and conjugates were eluted in 2x Laemmli buffer supplemented with 5 mM DTT at 70° C for 10 min prior to SDS-PAGE (Bio-Rad, 456–1094). Immunoprecipitation for exogenous TRAF6, ECSIT, NRDP1, and Parkin were performed using FLAG clonal M2 (1:1000; Sigma, Catalog# F104), HA (1:1000, Santa Cruz, Catalog#sc-805), and myc-HRP (1:1000; Roche, Catalog#a790-4628) antibodies, respectively. For FLAG and HA antibodies, secondary anti-mouse light chain antibody (Jackson Immunoresearch) was used.

### Complex I respiration assessments

Islets were isolated from mice and incubated overnight at 37° C. The next day, islets were picked and washed twice with 1x PBS. The PBS was decanted, and islets were flash-frozen in liquid nitrogen. Islet samples were freeze-thawed three times, and then 10 μg of sample was loaded per well into a Seahorse XF96 microplate in 20 μL of MAS buffer (70 mM sucrose, 220 mM mannitol, 5 mM KH_2_PO_4_, 5 mM MgCl_2_, 1 mM EGTA, 2 mM HEPES pH 7.4). To assess complex I respiration, islets were supplied with NADH and then oxygen consumption rate (OCR) was calculated using the Seahorse XF96 Extracellular Flux analyzer, as described previously^111^. Data were normalized to mitochondrial content.

### Mitochondrial fractionation

sgScram or sgTraf6 Min6 cells were treated with either BSA control or 0.5 mM palmitate for 48 h. Cells were trypsinized and collected, then washed twice with ice cold 1x PBS. Cells were then resuspended in 300 μL of ice cold HIM buffer (220 mM mannitol, 70mM sucrose, 1mM EGTA, 10mM HEPES pH 7.5) supplemented with 0.2% BSA and 1x protease inhibitors and incubated on ice for 30 min. Cells were homogenized in a Dounce homogenizer on ice for 50 passes three times, separated by 20 min centrifugation at 600 x *g*, 4°C. The supernatants from each 600 x g spin were pooled and spun for 15 min at 8,000 x *g*, 4 °C.

The pellet was resuspended in 100 μL HIM buffer (without BSA). Mitochondrial lysates were quantified via BCA assay (Pierce).

### TMT labeling mass spectrometry (MS)-based quantitative proteomics

#### Preparation of mitochondrial fractions

Mitochondrial fractions from sgScramble or sgTraf6 Min6 cells were collected as described above and were lysed in RIPA buffer (Thermofisher) containing phosphatase inhibitors. Mitochondrial lysates were quantified via BCA assay. All samples were diluted to 1 mg/mL in RIPA buffer.

#### Protein Digestion and TMT labeling

40 μL of each sample was submitted to the Proteomics Resource Facility at the University of Michigan for processing and mass spectrometry data acquisition. Briefly, upon reduction (5 mM DTT, for 30 min at 45 °C) and alkylation (15 mM 2-chloroacetamide, for 30 min at room temperature) of cysteines in samples, the proteins were precipitated by adding 6 volumes of ice-cold acetone followed by overnight incubation at-20 °C. The precipitate was spun down, and the pellet was allowed to air dry. The pellet was resuspended in 0.1M TEAB and overnight (∼16 h) digestion with trypsin/Lys-C mix (1:25 protease:protein; Promega) at 37 °C was performed with constant mixing using a thermomixer. The TMT 6-plex reagents were dissolved in 41 μL of anhydrous acetonitrile and labeling was performed by transferring the entire digest to TMT reagent vial and incubating at room temperature for 1 h. Reaction was quenched by adding 8 μL of 5% hydroxyl amine and further 15 min incubation. Labeled samples were mixed, and dried using a vacufuge. An offline fractionation of the combined sample (∼200 μg) into 8 fractions was performed using high pH reversed-phase peptide fractionation kit according to the manufacturer’s protocol (Pierce; Cat #84868). Fractions were dried and reconstituted in 9 μL of 0.1% formic acid/2% acetonitrile in preparation for LC-MS/MS analysis. Samples sgScram BSA 1-4 were labeled with TMT channels 126, 127N, 128N, and 129N, respectively. Samples sgTraf6 BSA 1-4 were labeled with TMT channels 130N, 131N, 132N, and 133N, respectively. Samples sgScram palm 1-4 were labeled with TMT channels 134N, 127C, 128C, and 129C, respectively. Samples sgTraf6 palm 1-4 were labeled with TMT channels 130C, 131C, 132C, and 133C, respectively.

#### Liquid chromatography-mass spectrometry analysis (LC-multinotch MS3)

To obtain superior quantitation accuracy, we employed multinotch-MS3 which minimizes the reporter ion ratio distortion resulting from fragmentation of co-isolated peptides during MS analysis^112^. Orbitrap Ascend Tribrid equipped with FAIMS source (Thermo Fisher Scientific) and Vanquish Neo UHPLC was used to acquire the data. 2 μL of the sample was resolved on an Easy-Spray PepMap Neo column (75 μm i.d. x 50 cm; Thermo Scientific) at the flow-rate of 300 nL/min using 0.1% formic acid/acetonitrile gradient system (3-19% acetonitrile in 72 min;19--29% acetonitrile in 28 min; 29-41% in 20 min followed by 10 min column wash at 95% acetonitrile and re-equilibration) and directly sprayed onto the mass spectrometer using EasySpray source (Thermo Fisher Scientific). FAIMS source was operated in standard resolution mode, with a nitrogen gas flow of 4.2 L/min, and inner and outer electrode temperature of 100 C and dispersion voltage or-5000 V. Two compensation voltages (CVs) of-45 and-65 V, 1.5 seconds per CV, were employed to select ions that enter the mass spectrometer for MS1 scan and MS/MS cycles. Mass spectrometer was set to collect MS1 scan (Orbitrap; 400-1600 m/z; 120K resolution; AGC target of 100%; max IT in Auto) following which precursor ions with charge states of 2-6 were isolated by quadrupole mass filter at 0.7 m/z width and fragmented by collision induced dissociation in ion trap (NCE 30%; normalized AGC target of 100%; max IT 35 ms). For multinotch-MS3, top 10 precursors from each MS2 were fragmented by HCD followed by Orbitrap analysis (NCE 55; 45K resolution; normalized AGC target of 200%; max IT 200 ms, 100-500 m/z scan range).

#### Data analysis

Proteome Discoverer (v3.0; Thermo Fisher) was used for data analysis. MS2 spectra were searched against SwissProt mouse protein database (Mus musculus (sp_canonical TaxID=10090) (v2023-09-13), downloaded on 05/15/2023) using the following search parameters: MS1 and MS2 tolerance were set to 10 ppm and 0.6 Da, respectively; carbamidomethylation of cysteines (57.02146 Da) and TMT labeling of lysine and N-termini of peptides (229.16293 Da) were considered static modifications; oxidation of methionine (15.9949 Da) and deamidation of asparagine and glutamine (0.98401 Da) were considered variable. Identified proteins and peptides were filtered to retain only those that passed ≤1% FDR threshold. Quantitation was performed using high-quality MS3 spectra (Average signal-to-noise ratio of 8 and <60% isolation interference).

### Statistics

In all figures, data are presented as means ± SEM, and error bars denote SEM, unless otherwise noted in the legends. Outlier tests (ROUT method^113^) were routinely performed in GraphPad Prism. Statistical comparisons were performed using unpaired two-tailed Student’s *t*-tests, 1-way or 2-way ANOVA, followed by Tukey’s or Sidak’s post-hoc test for multiple comparisons, as appropriate (GraphPad Prism). A *P* value < 0.05 was considered significant.

## Data Availability

Proteomics data deposition at ProteomeXchange is in process and will be publicly available at the date of publication. Microscopy data will be shared by the lead contact on request. RNAseq data have been deposited at GEO and will be publicly available at the date of publication.

Any additional information required to reanalyze the data reported in this paper is available from the corresponding author on request.

## ACKNOWLEDGEMENTS

Next generation sequencing was performed in the Advanced Genomics Core at the University of Michigan. Proteomics studies were performed in the Proteomics Resource Facility at the University of Michigan. Human pancreatic islets were provided by the Alberta Diabetes Institute IsletCore at the University of Alberta in Edmonton (http://www.bcell.org/adi-isletcore.html) with the assistance of the Human Organ Procurement and Exchange (HOPE) program, Trillium Gift of Life Network (TGLN), and other Canadian organ procurement organizations. Islet isolation was approved by the Human Research Ethics Board at the University of Alberta (Pro00013094). All donors’ families gave informed consent for the use of pancreatic tissue in research. We thank Dr. K. Claiborn and members of the Soleimanpour laboratory for helpful advice.

## AUTHOR CONTRIBUTIONS

E.L-D. conceived, designed, and performed experiments, interpreted results, drafted, and reviewed the manuscript. E.M.W, J.Z., Y.D., V.S., A.M.S., E.C.R., B.A.H-K., A.C.L, M.B.P., D.L.H., V.B., and L.S. designed and performed experiments and interpreted results. S.G., A.I.N., and O.S.S. designed studies, interpreted results, and reviewed the manuscript. S.A.S. conceived and designed the studies, interpreted results, drafted, edited, and reviewed the manuscript.

