## Supplementary Material 1-2025 for "TRAF6 integrates innate immune signals to regulate glucose homeostasis via Parkin-dependent and-independent mitophagy"

**Corresponding Author**

Figure S1

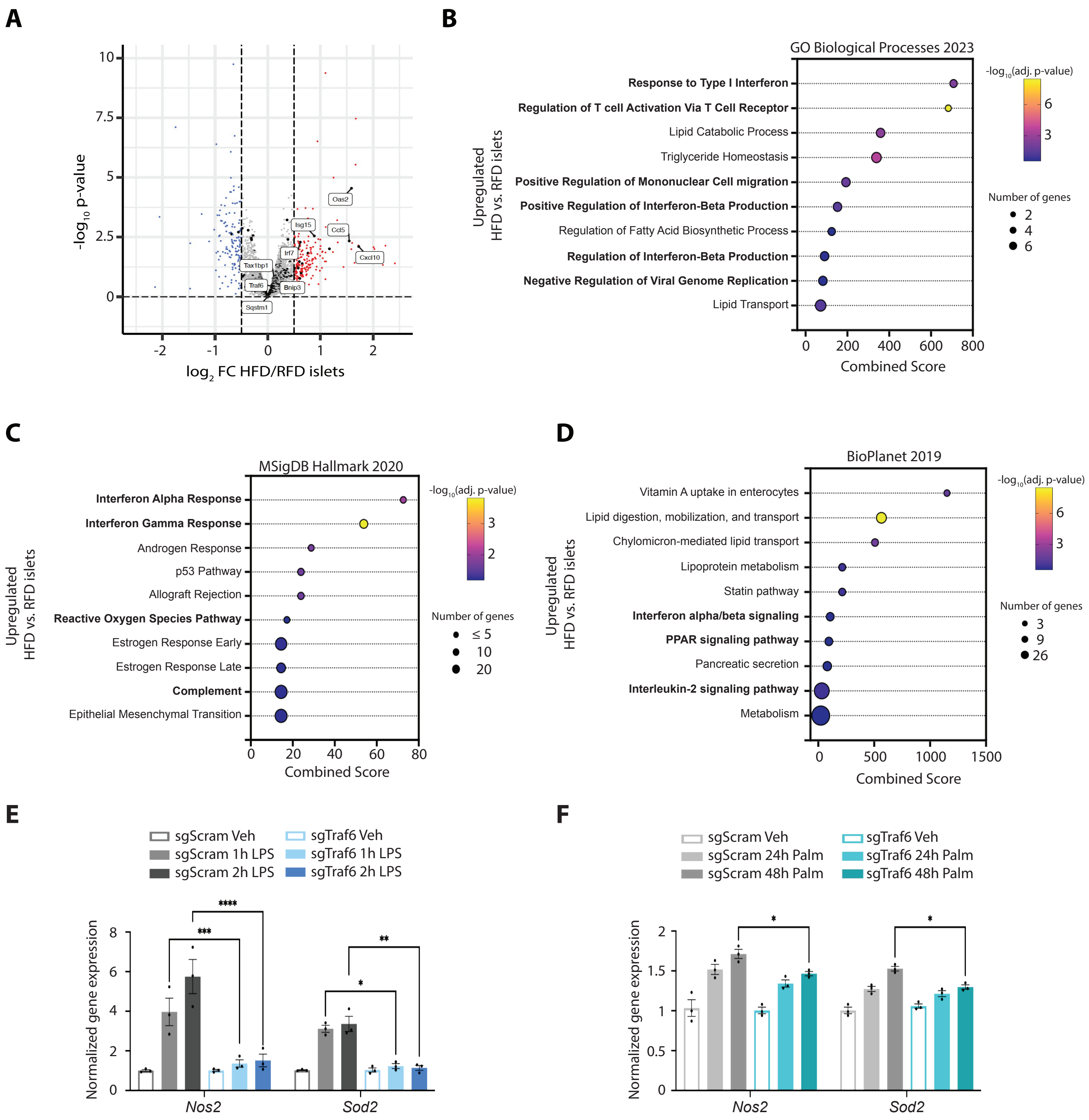
A
log2 FC HFD/RFD islets
-log10 p-value
Oas2
Isg15
Ccl5
Cxcl10
Irf7
Tax1bp1
Traf6
Bnip3
Sqstm1
B
GO Biological Processes 2023
Response to Type I Interferon
Regulation of T cell Activation Via T Cell Receptor
Lipid Catabolic Process
Triglyceride Homeostasis
Positive Regulation of Mononuclear Cell migration
Positive Regulation of Interferon-Beta Production
Regulation of Fatty Acid Biosynthetic Process
Regulation of Interferon-Beta Production
Negative Regulation of Viral Genome Replication
Lipid Transport
Upregulated HFD vs. RFD islets
Combined Score
-log10(adj. p-value)
Number of genes
C
MSigDB Hallmark 2020
Interferon Alpha Response
Interferon Gamma Response
Androgen Response
p53 Pathway
Allograft Rejection
Reactive Oxygen Species Pathway
Estrogen Response Early
Estrogen Response Late
Complement
Epithelial Mesenchymal Transition
Upregulated HFD vs. RFD islets
Combined Score
-log10(adj. p-value)
Number of genes
D
BioPlanet 2019
Vitamin A uptake in enterocytes
Lipid digestion, mobilization, and transport
Chylomicron-mediated lipid transport
Lipoprotein metabolism
Statin pathway
Interferon alpha/beta signaling
PPAR signaling pathway
Pancreatic secretion
Interleukin-2 signaling pathway
Metabolism
Upregulated HFD vs. RFD islets
Combined Score
-log10(adj. p-value)
Number of genes
E
sgScram Veh
sgScram 1h LPS
sgScram 2h LPS
sgTraf6 Veh
sgTraf6 1h LPS
sgTraf6 2h LPS
Normalized gene expression
Nos2
Sod2
F
sgScram Veh
sgScram 24h Palm
sgScram 48h Palm
sgTraf6 Veh
sgTraf6 24h Palm
sgTraf6 48h Palm
Normalized gene expression
Nos2
Sod2

**Figure S1. Metabolic stress elicits innate immune signaling pathways.**

**(A)** Volcano plot depicting differential RNA expression in islets isolated from control mice fed 12 weeks of HFD vs. control mice fed 12 weeks of RFD. *n* = 4-5 samples per group. Differentially expressed genes identified by *P* value < 0.05 and $|\log_2 FC| > 0.5$. **(B)** Selected terms from Enrichr analysis of differentially expressed upregulated genes in islets from from control mice fed 12 weeks of HFD vs. control mice fed 12 weeks of RFD analyzed with the following libraries: GO Biological Processes 2023, **(C)** MSigDB Hallmark 2020, and **(D)** BioPlanet 2019. **(E)** qPCR analysis of *Nos2* and *Sod2* in sgScram or sgTraf6 Min6 cells following exposure to vehicle (DMSO) or LPS (5 nM for 1 or 2 h). *n* = 3/group. **P* < 0.01, ***P* < 0.01, ****P* < 0.001, *****P* < 0.0001 by 2-way ANOVA. **(F)** qPCR analysis of *Nos2* and *Sod2* in sgScram or sgTraf6 Min6 cells following exposure to vehicle (BSA) or palmitate (0.5 mM for 24 or 48 h). *n* = 3/group. **P* < 0.05 by 2-way ANOVA.

Figure S2

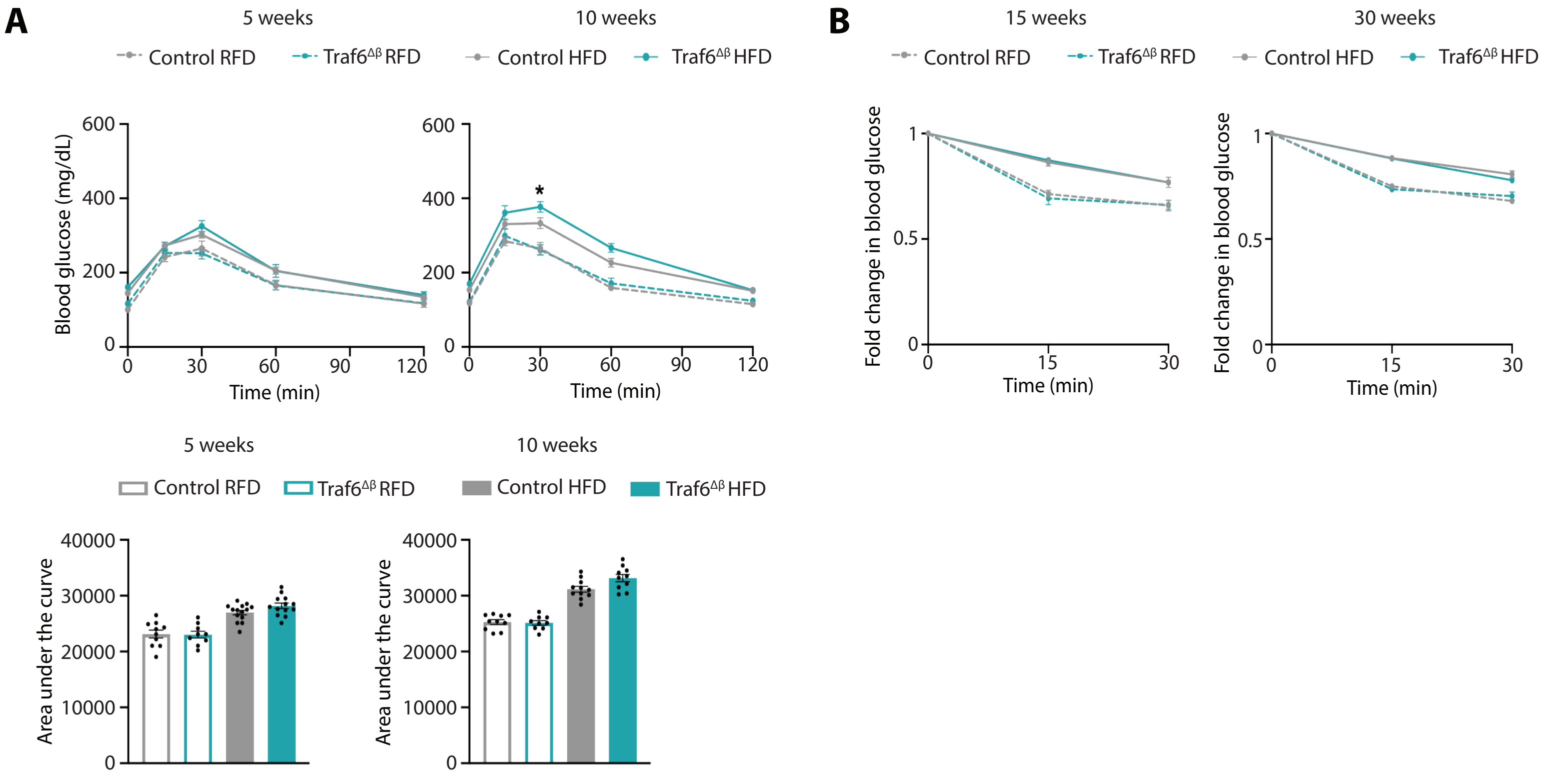
A
5 weeks
10 weeks
Control RFD
Traf6ΔB RFD
Control HFD
Traf6ΔB HFD
Blood glucose (mg/dL)
Time (min)
*
Area under the curve
B
15 weeks
30 weeks
Fold change in blood glucose

**Figure S2. TRAF6-deficiency results in progressive glucose intolerance in HFD-fed male mice.**

**(A)** Blood glucose concentrations measured during IPGTT (top) and AUC (bottom) of control and Traf6$^{\Delta\beta}$ male mice fed either RFD or HFD for 5 weeks (left) or 10 weeks (right). *n* = 9-14/group. *$P$ < 0.05 by 2-way ANOVA (IPGTT) or 1-way ANOVA (AUC). **(B)** Fold changes in blood glucose measured during *in vivo* insulin tolerance test in control and Traf6$^{\Delta\beta}$ male mice RFD or HFD for 15 weeks (left) or 30 weeks (right). *n* = 5-7/group.

Figure S3

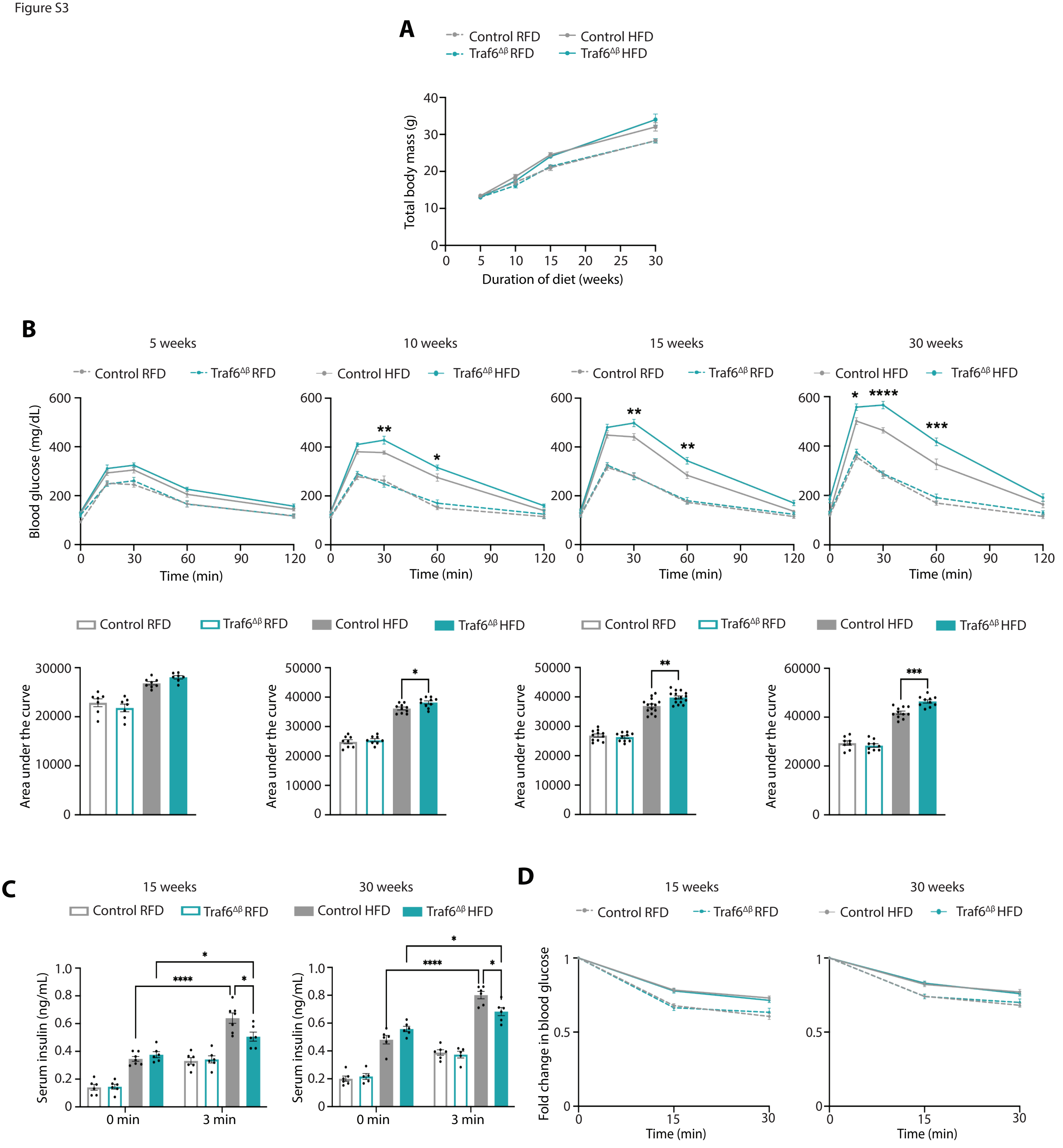
A
Control RFD
Control HFD
Traf6ΔB RFD
Traf6ΔB HFD
Total body mass (g)
Duration of diet (weeks)
B
5 weeks
10 weeks
15 weeks
30 weeks
Control RFD
Traf6ΔB RFD
Control HFD
Traf6ΔB HFD
Blood glucose (mg/dL)
Time (min)
Area under the curve
C
15 weeks
30 weeks
Serum insulin (ng/mL)
0 min
3 min
D
15 weeks
30 weeks
Fold change in blood glucose
Time (min)

**Figure S3. TRAF6-deficiency results in progressive glucose intolerance in HFD-fed female mice.**

**(A)** Total body mass of control or $Traf6^{\Delta\beta}$ female mice following 30 weeks of RFD or HFD. *n* = 7-10/group. **(B)** Blood glucose concentrations (top) and AUC (bottom) measured during IPGTT of control and $Traf6^{\Delta\beta}$ female mice fed RFD or HFD for 5, 10, 15, or 30 weeks, respectively. *n* = 7-15/group. *$P < 0.05$, **$P < 0.01$, ***$P < 0.001$, ****$P < 0.0001$ by 2-way ANOVA (IPGTT) or 1-way ANOVA (AUC). **(C)** Serum insulin levels measured during *in vivo* glucose-stimulated insulin release assays in control and $Traf6^{\Delta\beta}$ female mice fed RFD or HFD for 15 weeks (left) or 30 weeks (right). *n* = 5-7/group. *$P < 0.05$, ****$P < 0.0001$ by 1-way ANOVA. **(D)** Fold changes in blood glucose measured during *in vivo* insulin tolerance test in control and $Traf6^{\Delta\beta}$ female mice fed RFD or HFD for 15 weeks (left) or 30 weeks (right). *n* = 5-8/group.

Figure S4

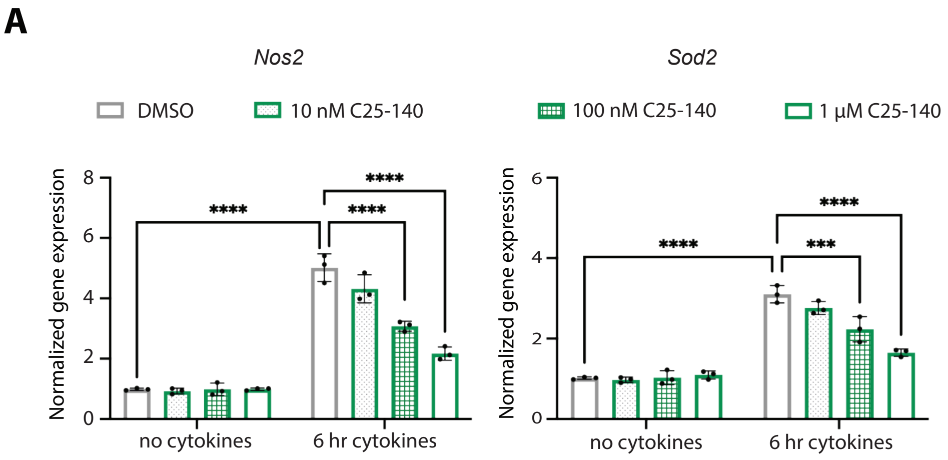
A
Nos2
Sod2
DMSO
10 nM C25-140
100 nM C25-140
1 µM C25-140
Normalized gene expression
8
6
4
2
0
****
****
****
no cytokines
6 hr cytokines
Normalized gene expression
6
4
2
0
****
****
***
no cytokines
6 hr cytokines

**Figure S4. The TRAF6 inhibitor C25-140 reduces NF-κB target gene expression in a dose-dependent manner.**

qPCR analysis of *Nos2* (left) and *Sod2* (right) in isolated islets from control mice fed RFD for 15 weeks following exposure to vehicle (DMSO), or 10 nM, 100 nM, or 1 μM C25-140 and cytokines (75 U/mL IL-1β, 750 U/mL TNF-α, and 750 U/mL IFN-γ) for 6 h. $n$ = 3/group. *$P$ < 0.05, **$P$ < 0.01, ***$P$ < 0.001, ****$P$ < 0.0001 by 2-way ANOVA.

Figure S5

A

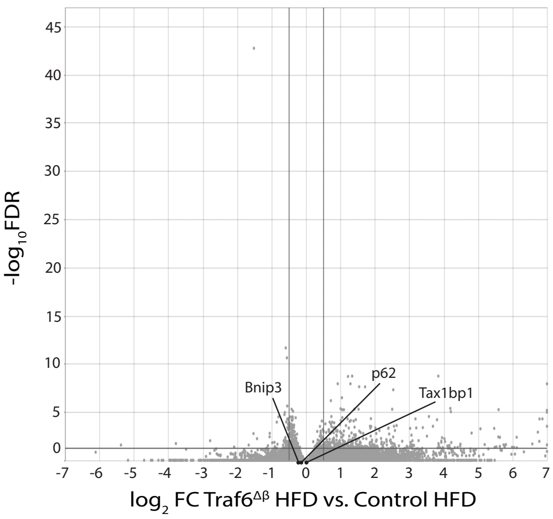

-log10FDR
45
40
35
30
25
20
15
10
5
0
Bnip3
p62
Tax1bp1
-7 -6 -5 -4 -3 -2 -1 0 1 2 3 4 5 6 7
log2 FC Traf6ΔB HFD vs. Control HFD


B

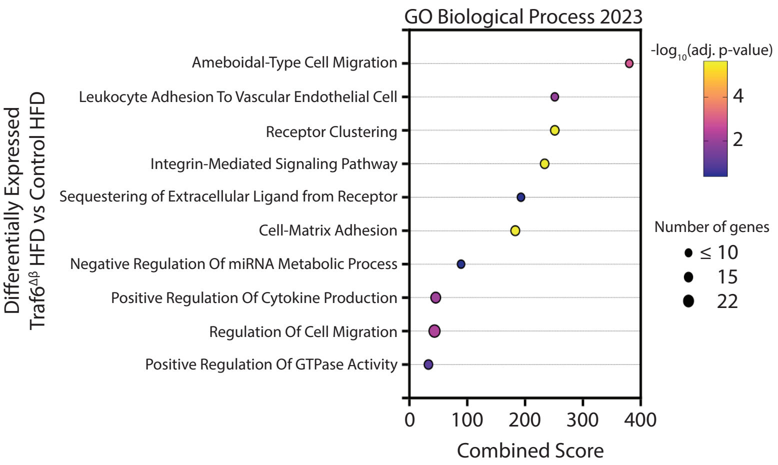

GO Biological Process 2023
Differentially Expressed
Traf6ΔB HFD vs Control HFD
Ameboidal-Type Cell Migration
Leukocyte Adhesion To Vascular Endothelial Cell
Receptor Clustering
Integrin-Mediated Signaling Pathway
Sequestering of Extracellular Ligand from Receptor
Cell-Matrix Adhesion
Negative Regulation Of miRNA Metabolic Process
Positive Regulation Of Cytokine Production
Regulation Of Cell Migration
Positive Regulation Of GTPase Activity
0 100 200 300 400
Combined Score
-log10(adj. p-value)
4
2
Number of genes
≤ 10
15
22

**Figure S5. Differential expression and pathway analysis of RNA-seq from islets of HFD-fed Traf6$^{\Delta\beta}$ mice.**

**(A)** Volcano plot depicting differential RNA expression in islets from Traf6$^{\Delta\beta}$ vs. control mice fed HFD for 15 weeks. *n* = 5-6/group. Significantly differentially expressed genes demarcated by FDR > 0.5 and $|\log_2 FC| > 0.5$. **(B)** Selected terms from Enrichr analysis of differentially expressed genes in islets in islets from Traf6$^{\Delta\beta}$ vs. control mice fed HFD for 15 weeks analyzed with the GO Biological Processes 2023 library.

Figure S6

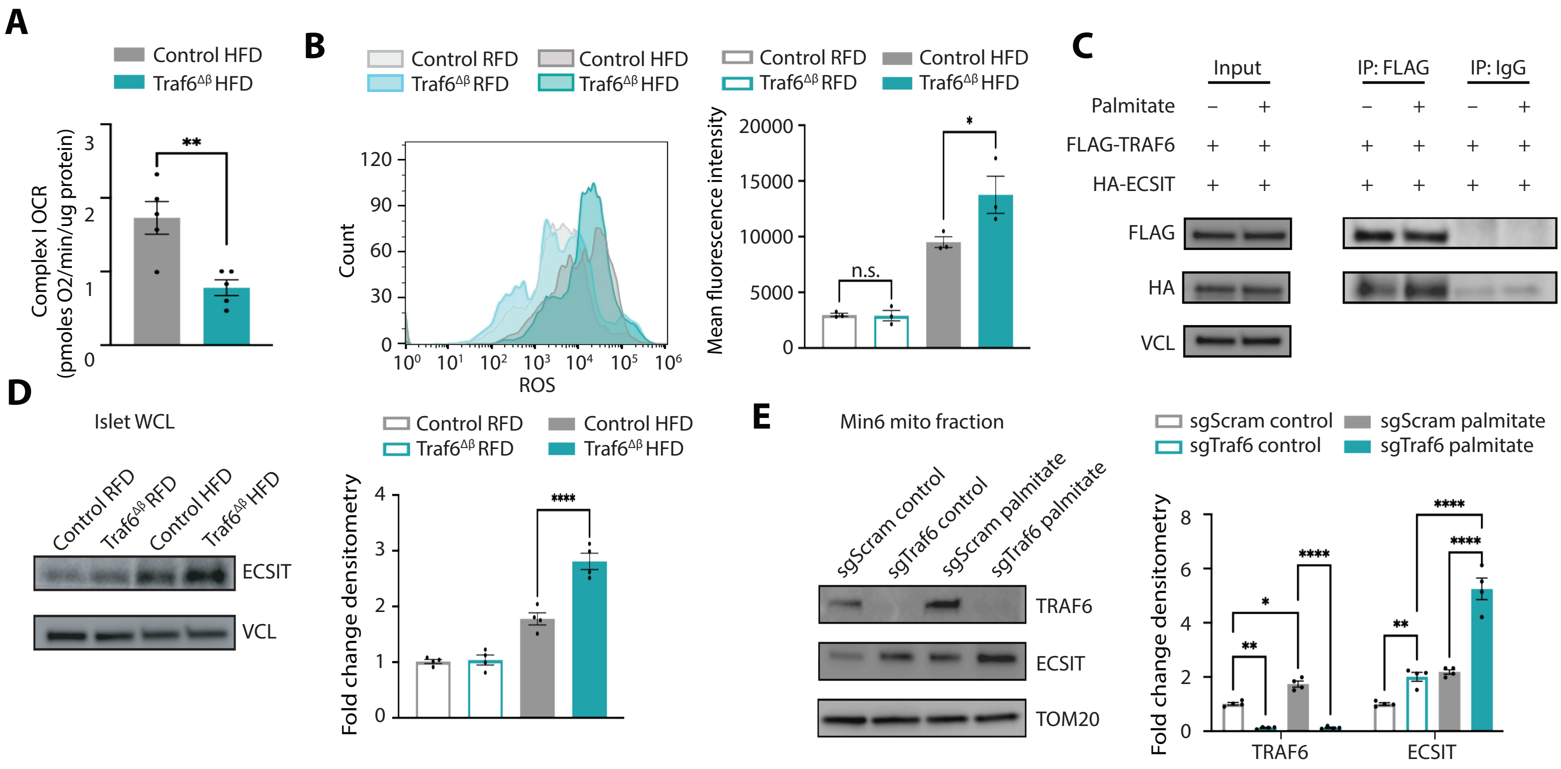
A
Control HFD
Traf6^Δβ HFD
Complex I OCR (pmoles O2/min/ug protein)
**
B
Control RFD
Control HFD
Traf6^Δβ RFD
Traf6^Δβ HFD
Count
ROS
Mean fluorescence intensity
n.s.
*
C
Input
IP: FLAG
IP: IgG
Palmitate
FLAG-TRAF6
HA-ECSIT
FLAG
HA
VCL
D
Islet WCL
Control RFD
Traf6^Δβ RFD
Control HFD
Traf6^Δβ HFD
ECSIT
VCL
Fold change densitometry
****
E
Min6 mito fraction
sgScram control
sgTraf6 control
sgScram palmitate
sgTraf6 palmitate
TRAF6
ECSIT
TOM20
Fold change densitometry

**Figure S6. TRAF6-deficiency increases expression and mitochondrial localization of ECSIT following metabolic stress.**

**(A)** Complex I respiration measured in islets isolated from control and Traf6$^{\Delta\beta}$ mice fed HFD for 15 weeks. *n* = 5 per group. $^{**}P < 0.01$ by unpaired Student's two-tailed t-test. **(B)** Representative ROS mean fluorescence intensity measured via flow cytometry (left) and quantification (right) in isolated islets from control and Traf6$^{\Delta\beta}$ mice fed RFD or HFD for 15 weeks. *n* = 3/ group. $^{*}P < 0.05$ by 1-way ANOVA. **(C)** Representative WB of FLAG and HA levels following anti-FLAG or anti-IgG immunoprecipitation (IP) in Min6 cells transfected with plasmids expressing FLAG-TRAF6 and HA-ECSIT and exposed to BSA or 0.5 mM palmitate for 48 h. VCL served as a loading control. *n* = 3/group. **(D)** Protein expression and densitometry quantification (relative to Control RFD) of ECSIT by WB in islets isolated from control or Traf6$^{\Delta\beta}$ mice fed RFD or HFD for 15 weeks. VCL served as a loading control. *n* = 4/group. $^{****}P < 0.0001$ by 1-way ANOVA. **(E)** Protein expression and densitometry quantification (relative to sgScram control) of TRAF6 and ECSIT by WB in mitochondrial fractions from sgScram and sgTraf6 Min6 cells exposed to BSA or 0.5 mM palmitate for 48 h. TOM20 served as a loading control. *n* = 4/group. $^{*}P < 0.05$, $^{**}P < 0.01$, $^{****}P < 0.0001$ by 2-way ANOVA.

Figure S7

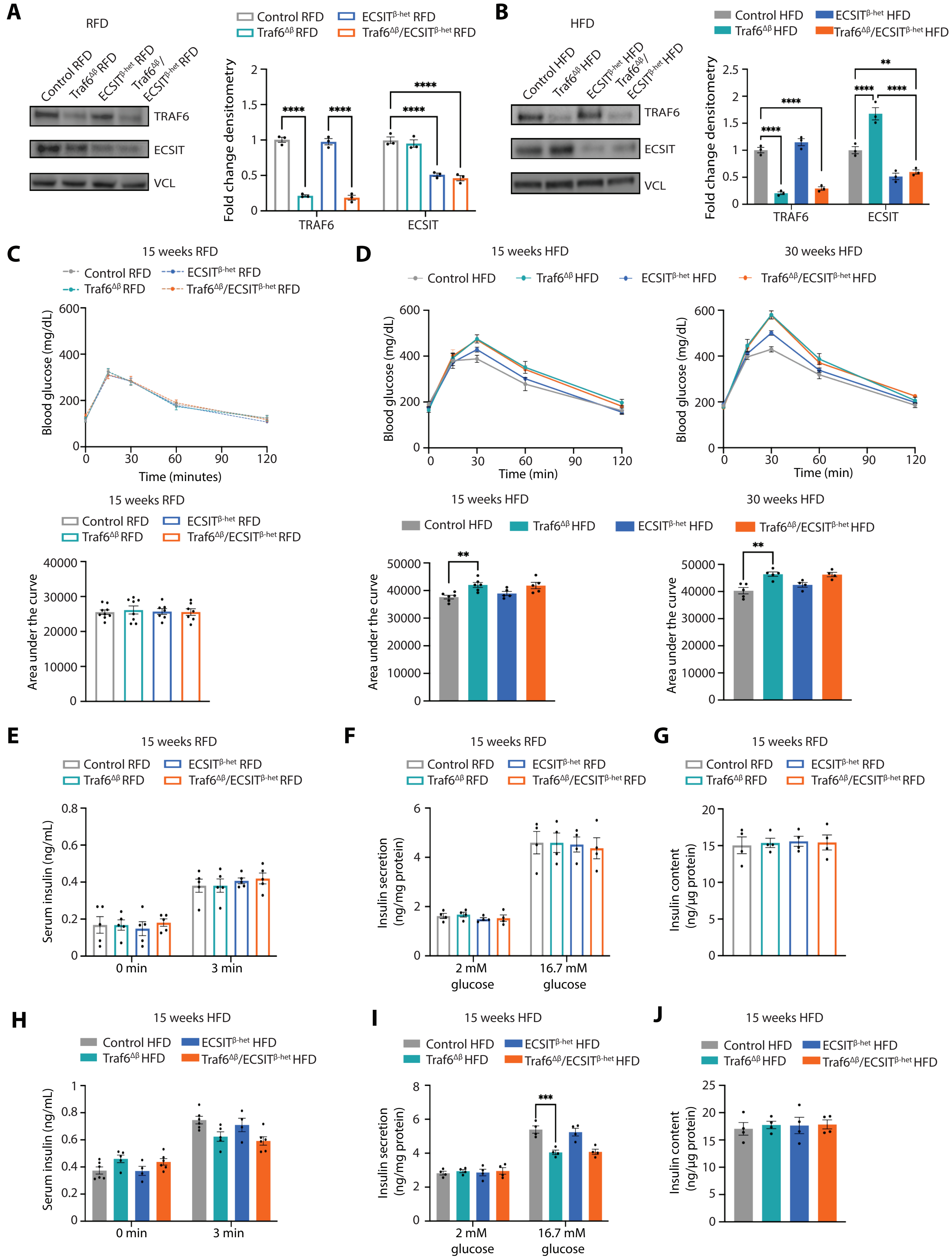
A
RFD
Control RFD
Traf6ΔB RFD
ECSITB-het RFD
Traf6ΔB/ECSITB-het RFD
TRAF6
ECSIT
VCL
Fold change densitometry
****
TRAF6
ECSIT
B
HFD
Control HFD
Traf6ΔB HFD
ECSITB-het HFD
Traf6ΔB/ECSITB-het HFD
TRAF6
ECSIT
VCL
Fold change densitometry
****
**
C
15 weeks RFD
Blood glucose (mg/dL)
Time (minutes)
Area under the curve
D
15 weeks HFD
30 weeks HFD
Blood glucose (mg/dL)
Time (min)
Area under the curve
**
E
15 weeks RFD
Serum insulin (ng/mL)
0 min
3 min
F
15 weeks RFD
Insulin secretion (ng/mg protein)
2 mM glucose
16.7 mM glucose
G
15 weeks RFD
Insulin content (ng/µg protein)
H
15 weeks HFD
Serum insulin (ng/mL)
0 min
3 min
I
15 weeks HFD
Insulin secretion (ng/mg protein)
***
2 mM glucose
16.7 mM glucose
J
15 weeks HFD
Insulin content (ng/µg protein)

**Figure S7. Increases in ECSIT expression do not elicit β-cell dysfunction in HFD-fed Traf6$^{\Delta\beta}$ mice.**

**(A)** Protein expression and densitometry quantification (relative to Control RFD) of ECSIT by WB in islets isolated from control, Traf6$^{\Delta\beta}$, ECSIT$^{\beta\text{-het}}$, or Traf6$^{\Delta\beta}$/ECSIT$^{\beta\text{-het}}$ mice fed RFD for 15 weeks. VCL served as a loading control. *n* = 3/group. *****P* < 0.0001 by 2-way ANOVA. **(B)** Protein expression and densitometry quantification (relative to Control RFD) of ECSIT by WB in islets isolated from control, Traf6$^{\Delta\beta}$, ECSIT$^{\beta\text{-het}}$, or Traf6$^{\Delta\beta}$/ECSIT$^{\beta\text{-het}}$ mice fed HFD for 15 weeks. VCL served as a loading control. *n* = 3/group. ***P* < 0.01, *****P* < 0.0001 by 2-way ANOVA. **(C)** Blood glucose concentrations (top) and AUC (bottom) measured during IPGTT of control, Traf6$^{\Delta\beta}$, ECSIT$^{\beta\text{-het}}$, or Traf6$^{\Delta\beta}$/ECSIT$^{\beta\text{-het}}$ male mice fed RFD for 15 weeks. *n* = 7-9/group. **(D)** Blood glucose concentrations (top) and AUC (bottom) measured during IPGTT of control, Traf6$^{\Delta\beta}$, ECSIT$^{\beta\text{-het}}$, or Traf6$^{\Delta\beta}$/ECSIT$^{\beta\text{-het}}$ male mice fed HFD for 15 weeks (left) or 30 weeks (right) HFD. *n* = 4-6/group. ***P* < 0.01 by 1-way ANOVA (AUC). **(E)** Serum insulin levels measured during *in vivo* glucose-stimulated insulin release assays in control, Traf6$^{\Delta\beta}$, ECSIT$^{\beta\text{-het}}$, or Traf6$^{\Delta\beta}$/ECSIT$^{\beta\text{-het}}$ male mice fed RFD for 15 weeks. *n* = 5-7/group. **(F)** Glucose-stimulated insulin secretion from islets isolated from control, Traf6$^{\Delta\beta}$, ECSIT$^{\beta\text{-het}}$, or Traf6$^{\Delta\beta}$/ECSIT$^{\beta\text{-het}}$ mice fed RFD for 15 weeks. *n*=4/group. **(G)** Insulin content in islets from control, Traf6$^{\Delta\beta}$, ECSIT$^{\beta\text{-het}}$, or Traf6$^{\Delta\beta}$/ECSIT$^{\beta\text{-het}}$ mice fed RFD for 15 weeks. *n* = 4/group. **(H)** Serum insulin levels measured during *in vivo* glucose-stimulated insulin release assays in control, Traf6$^{\Delta\beta}$, ECSIT$^{\beta\text{-het}}$, or Traf6$^{\Delta\beta}$/ECSIT$^{\beta\text{-het}}$ male mice fed HFD for 15 weeks. *n* = 4-6/ group. **(I)** Glucose-stimulated insulin secretion from islets isolated from control, Traf6$^{\Delta\beta}$, ECSIT$^{\beta\text{-het}}$, or Traf6$^{\Delta\beta}$/ECSIT$^{\beta\text{-het}}$ mice fed HFD for 15 weeks. *n* = 4/group. ****P* < 0.001 by 2-way ANOVA. **(J)** Insulin content in islets from control, Traf6$^{\Delta\beta}$, ECSIT$^{\beta\text{-het}}$, or Traf6$^{\Delta\beta}$/ECSIT$^{\beta\text{-het}}$ mice fed HFD for 15 weeks. *n* = 4/group.

Figure S8

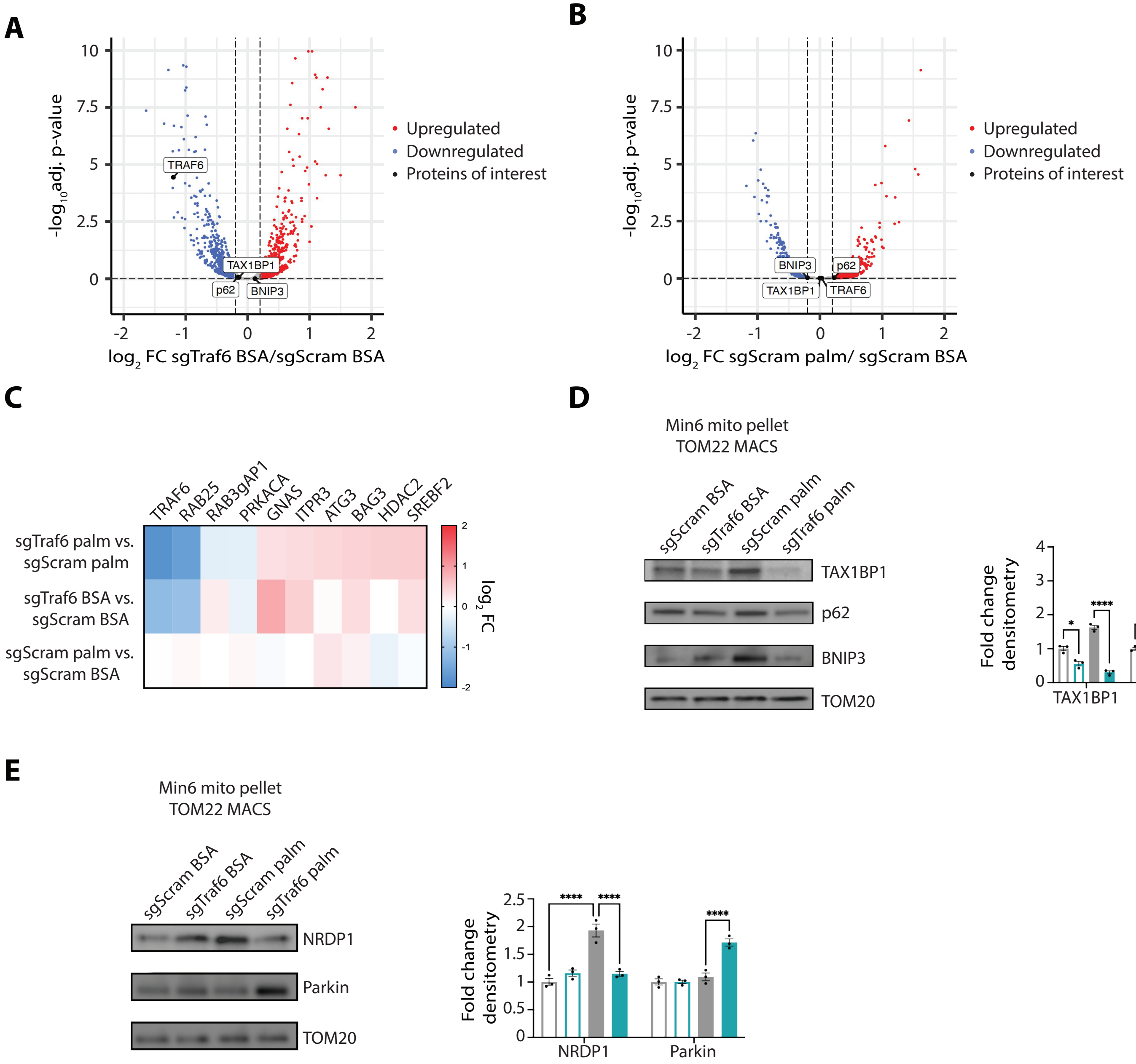

A
B
C
D
E
Upregulated
Downregulated
Proteins of interest
-log10adj. p-value
log2 FC sgTraf6 BSA/sgScram BSA
log2 FC sgScram palm/ sgScram BSA
TRAF6
TAX1BP1
p62
BNIP3
TRAF6
RAB25
RAB3gAP1
PRKACA
GNAS
ITPR3
ATG3
BAG3
HDAC2
SREBF2
sgTraf6 palm vs. sgScram palm
sgTraf6 BSA vs. sgScram BSA
sgScram palm vs. sgScram BSA
log2 FC
Min6 mito pellet
TOM22 MACS
sgScram BSA
sgTraf6 BSA
sgScram palm
sgTraf6 palm
TAX1BP1
p62
BNIP3
TOM20
Fold change densitometry
NRDP1
Parkin
TOM20

**Figure S8. Palmitate alters recruitment of the mitophagy machinery in TRAF6-deficient β-cells.**

**(A)** Volcano plot of differentially expressed proteins in sgTraf6 control vs. sgScram control mitochondrial fractions. Differentially expressed proteins identified by adjusted *P* value < 0.05 and $|\log_2 FC| > 0.25$. *n* = 4/group. **(B)** Volcano plot of differentially expressed proteins in sgScram palm vs. sgScram control mitochondrial fractions. Differentially expressed proteins identified by adjusted *P* value < 0.05 and $|\log_2 FC| > 0.25$. *n* = 4/group. **(C)** Differential protein expression heatmap of the 10 unselected mitophagy and autophagy proteins identified in Figure 5C, displayed as integrated mean $\log_2 FC$, from proteomics studies of mitochondrial fractions of sgTraf6 vs. sgScram Min6 cells following exposure to 0.5 mM palmitate or BSA for 48 h. *n* = 4/group. **(D)** WB with densitometry of TAX1BP1, p62, and BNIP3 (relative to sgScram control) in mitochondrial fractions (isolated via TOM22 MACS kit) from sgScram and sgTraf6 Min6 cells exposed to BSA or 0.5 mM palmitate for 48 h. TOM20 served as a loading control. *n* = 3/group. **P* < 0.05, *****P* < 0.0001 by 2-way ANOVA. **(E)** WB with densitometry of NRDP1 and Parkin (relative to sgScram control) in mitochondrial fractions (isolated via TOM22 MACS kit) from sgScram and sgTraf6 Min6 cells exposed to BSA or 0.5 mM palmitate for 48 h. TOM20 served as a loading control. *n* = 3/group. *****P* < 0.0001 by 2-way ANOVA.

Figure S9

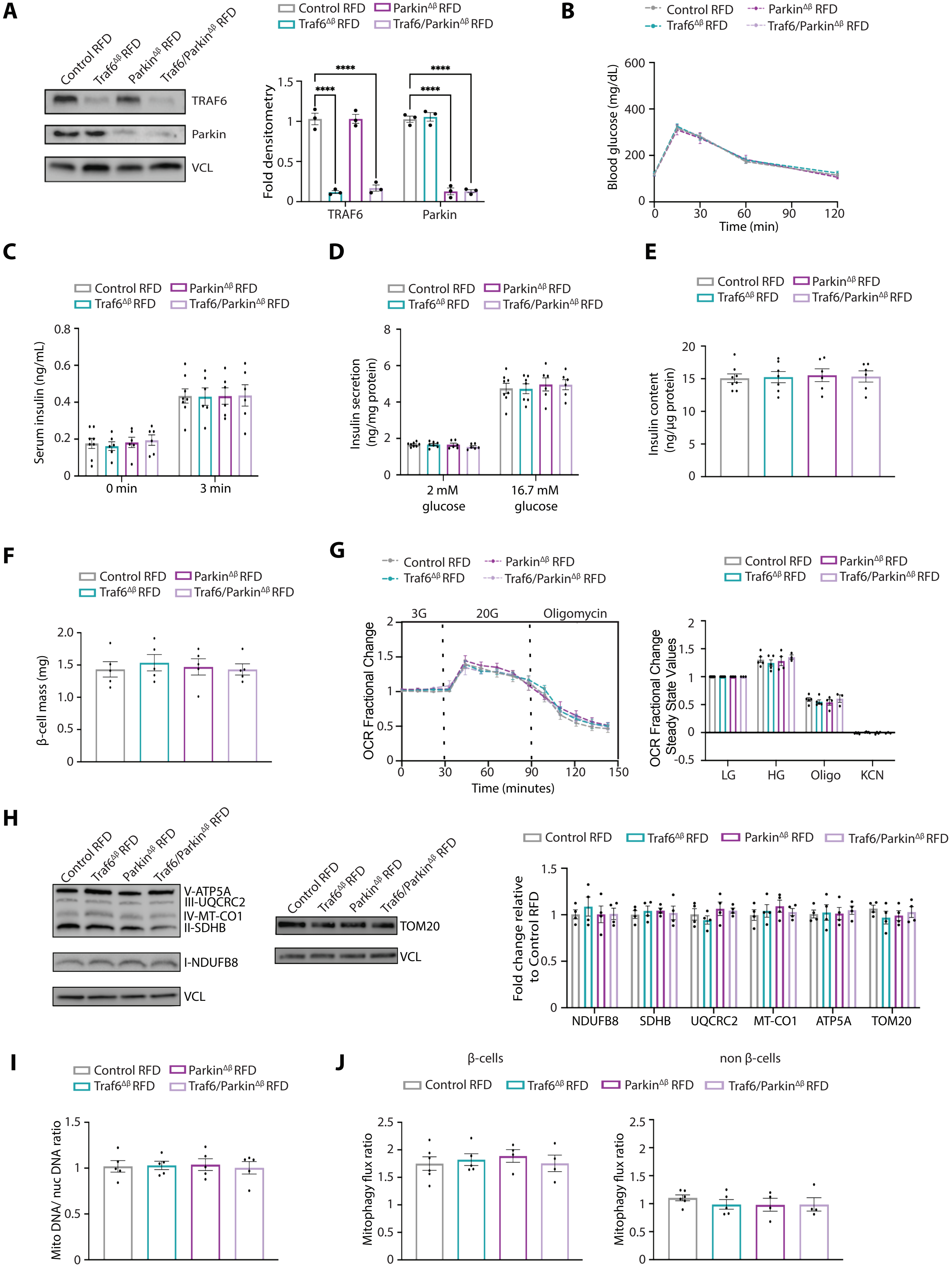
A
Control RFD
Traf6^Δβ RFD
Parkin^Δβ RFD
Traf6/Parkin^Δβ RFD
TRAF6
Parkin
VCL
Fold densitometry
****
TRAF6
Parkin
B
Blood glucose (mg/dL)
Time (min)
C
Serum insulin (ng/mL)
0 min
3 min
D
Insulin secretion (ng/mg protein)
2 mM glucose
16.7 mM glucose
E
Insulin content (ng/µg protein)
F
β-cell mass (mg)
G
3G
20G
Oligomycin
OCR Fractional Change
Time (minutes)
OCR Fractional Change Steady State Values
LG
HG
Oligo
KCN
H
V-ATP5A
III-UQCRC2
IV-MT-CO1
II-SDHB
I-NDUFB8
VCL
TOM20
Fold change relative to Control RFD
NDUFB8
SDHB
UQCRC2
MT-CO1
ATP5A
TOM20
I
Mito DNA/ nuc DNA ratio
J
β-cells
non β-cells
Mitophagy flux ratio

**Figure S9. Combined TRAF6- and Parkin-deficiency do not impact physiologic or mitochondrial function in RFD-fed mice.**

**(A)** Protein expression and densitometry quantification (relative to Control RFD) of TRAF6 and Parkin by WB in isolated islets from control, Traf6$^{\Delta\beta}$, Parkin$^{\Delta\beta}$, and Traf6/Parkin$^{\Delta\beta}$ mice fed RFD for 15 weeks. VCL served as a loading control. *n* = 3/group. ****$P$ < 0.0001 by 1-way ANOVA. **(B)** Blood glucose concentrations measured during IPGTT of control, Traf6$^{\Delta\beta}$, Parkin$^{\Delta\beta}$, and Traf6/Parkin$^{\Delta\beta}$ male mice fed RFD for 15 weeks. *n* = 8-11/group. **(C)** Serum insulin levels measured during *in vivo* glucose-stimulated insulin release assays in male mice fed RFD for 15 weeks. *n* = 6-8/group. **(D)** Glucose-stimulated insulin secretion from islets isolated from control, Traf6$^{\Delta\beta}$, Parkin$^{\Delta\beta}$, and Traf6/Parkin$^{\Delta\beta}$ mice fed RFD for 15 weeks. *n* = 6-8/group. **(E)** Insulin content in islets from control, Traf6$^{\Delta\beta}$, Parkin$^{\Delta\beta}$, and Traf6/Parkin$^{\Delta\beta}$ mice fed HFD for 15 weeks. *n* = 6-8/ group. **(F)** Pancreatic β-cell mass from control, Traf6$^{\Delta\beta}$, Parkin$^{\Delta\beta}$, and Traf6/Parkin$^{\Delta\beta}$ mice fed RFD for 15 weeks. *n* = 5/group. **(G)** Oxygen consumption rate (OCR) fractional change (left) and steady state quantification (right) following exposure to 3 mM glucose, 20 mM glucose, 10 μM oligomycin, and 3 mM KCN in isolated islets from control, Traf6$^{\Delta\beta}$, Parkin$^{\Delta\beta}$, and Traf6/Parkin$^{\Delta\beta}$ islets isolated from mice fed RFD for 15 weeks. *n* = 3-5/group. **(H)** WB for OXPHOS subunits and TOM20 with densitometry (relative to Control HFD) in islets from control, Traf6$^{\Delta\beta}$, Parkin$^{\Delta\beta}$, and Traf6/Parkin$^{\Delta\beta}$ mice fed RFD for 15 weeks. VCL served as a loading control. *n* = 4/group. **(I)** mtDNA to nuclear DNA ratio in isolated islets from control, Traf6$^{\Delta\beta}$, Parkin$^{\Delta\beta}$, and Traf6/Parkin$^{\Delta\beta}$ mice fed RFD for 15 weeks. *n* =4-5/group. **(J)** Flow cytometry quantification of mitophagy flux by MtPhagy dye approach in β-cells from control, Traf6$^{\Delta\beta}$, Parkin$^{\Delta\beta}$, and Traf6/Parkin$^{\Delta\beta}$ mice fed HFD for 15 weeks. *n* = 4-5/group.

**Table S1. Luciferase plasmids and pathways.**

| Plasmid name | Pathway | Transcription Factors |
|---|---|---|
| pLminP_Luc2P_RE1 | NFκB | NFKB1, RELA (p50, p65) |
| pLminP_Luc2P_RE38 | RELB | RELB |
| pLminP_Luc2P_RE10 | IRF3 | IRF3 |
| pLminP_Luc2P_RE5 | SIE (sis-inducible element) | STAT3 |
| pLminP_Luc2P_RE52 | STAT1 | STAT1 |
| pLminP_Luc2P_RE57 | Type I Interferon | STAT1, STAT2 |
| pLminP_Luc2P_RE27 | Type II Interferon | STAT1 |
| pLminP_Luc2P_RE47 | IFNB1 | AP-1, IRF3, IRF7, NFκB |
| pLminP_Luc2P_RE34 | MAPK | ELK1 |
| pLminP_Luc2P_RE28 | Lysosomal biogenesis | TFEB |
| pLminP_Luc2P_RE26 | C/EBPβ | CEBPB |
| pLminP_Luc2P_RE23 | Amino Acid Deprivation (AARE) | ATF4 |
| pLminP_Luc2P_RE21 | p53 | TRP53 |

**Table S2. Mitochondrial proteomics analysis of ubiquitin receptors and parkin-independent mitophagy receptors.**

| Protein Group | Protein name | sgTraf6 palm vs. sgScram palm | | sgTraf6 BSA vs. sgScram BSA | | sgScram palm vs. sgScram BSA | |
|---|---|---|---|---|---|---|---|
| | | log2FC | Adj. *P* value | log2FC | Adj. *P* value | log2FC | Adj. *P* value |
| Ubiquitin receptors | CALCOCO2 /NDP52 | ND | NA | ND | NA | ND | NA |
| | NBR1 | -0.03 | 0.57 | -0.05 | 0.99 | -0.28 | 0.99 |
| | OPTN | 0.08 | 0.98 | 0.16 | 0.99 | 0.11 | 0.98 |
| Mitophagy receptors | BCL2L13 | -0.09 | 0.96 | 0.13 | 0.83 | 0.02 | 0.99 |
| | NIX/BNIP3L | ND | NA | ND | NA | ND | NA |
| | FUNDC1 | -0.26 | 0.66 | -0.03 | 0.99 | -0.01 | 0.99 |

*ND= not detected, *NA= not applicable

**Table S3. Clinical characteristics of human islet donors.**

| Unique Islet Prep Identifier | Islet Isolation Center | Age (year) | BMI | Sex | HbA1c (%) | Cause of Death |
|---|---|---|---|---|---|---|
| HP-23200-01 | Prodo labs | 57 | 30.8 | M | 5.2 | Anorexic event |
| HP-23219-01 | Prodo labs | 59 | 24.5 | M | 5.2 | Stroke |
| R505 | ADI Islet Core | 53 | 33.1 | F | 5.4 | Undisclosed |
| HP-23240-01 | Prodo labs | 33 | 33.1 | M | 5.7 | Anorexic event |
| HP-23270-01 | Prodo labs | 34 | 27.0 | M | 5.5 | Stroke |
| HP-23301-01 | Prodo labs | 64 | 25.5 | M | 5.2 | Stroke |

**Table S4. Antibody information**

| Antibody | Company | Catalog Number |
|---|---|---|
| BNIP3 | Abcam | ab109362 |
| Cyclophilin B (PPIB) | ThermoFisher | PA1-027A |
| ECSIT | Abcam | ab21288 |
| FLAG clonal M2 | Sigma | F104 |
| HA | Santa Cruz | sc-805 |
| Insulin | Invitrogen | 701265 |
| Insulin | Fitzgerald | 20-IP35 |
| LC3 | Sigma | L-8919 |
| MFN2 | Abcam | ab56889 |
| myc-HRP | Roche | a790-4628 |
| NRDP1 | Santa Cruz | sc-365622 |
| Total OXPHOS | Abcam | ab110413 |
| p62 | Enzo | BML-PW9860 |
| Parkin | Cell Signaling | 4211 |
| Phospho-ubiquitin (Ser65) | Cell Signaling | 62802 |
| SDHA | Abcam | ab14715 |
| TAX1BP1 | Proteintech | 14424-1-AP |
| TOM20 | Cell Signaling | 42406 |
| TRAF6 | Abcam | ab40675 |
| Vinculin (VCL) | Millipore | CP74 |

**Table S5. Primer sequences**

| Primer | Forward | Reverse |
|---|---|---|
| *Hprt* | GGCCAGACTTTGTTGGATTTG | TGCGCTCATCTTAGGCTTTGT |
| *Mt9/11* | GAGCATCTTATCCACGCTTCC | GGTGGTACTCCCGCTGTAAA |
| *Nos2* | ACTGGGACAGCACAGAATGTTCC | CCAAATGTGCTTGTCACCACCAG |
| *Ndufv1* | CTTCCCCACTGGCCTCAAG | CCAAAACCCAGTGATCCAGC |
| *Sod2* | TACAACTCAGGTCGCTCTTCAGC | AGCCTCCAGCAACTCTCCTTT |
| *Traf6* | GCCTGTTTTCTTTTTACTCACCTG | CAAACCTCTACGGGTCATGC |
